# Panoramic visual representation in the dorsal visual pathway and its role in reorientation

**DOI:** 10.1101/827667

**Authors:** Tianyi Li, Angelo Arleo, Denis Sheynikhovich

## Abstract

While primates are primarily visual animals, how visual information is processed on its way to memory structures and contributes to the generation of visuospatial behaviors is poorly understood. Recent imaging data demonstrate the existence of scene-sensitive areas in the dorsal visual path that are likely to combine visual information from successive egocentric views, while behavioral evidence indicates the memory of surrounding visual space in extraretinal coordinates. The present work focuses on the computational nature of a panoramic representation that is proposed to link visual and mnemonic functions during natural behavior. In a spiking neural network model of the dorsal visual path it is shown how time-integration of spatial views can give rise to such a representation and how it can subsequently be used to perform memory-based spatial reorientation and visual search. More generally, the model predicts a common role of view-based allocentric memory storage in spatial and non-spatial mnemonic behaviors.

## Introduction

Recent breathtaking advances in our understanding of rodent hippocampal memory system pave the way for elucidating the organization of human spatial memory (*Burgess, 2014*; *Moser et al., 2017*). One major difference between primates and rodents is the role of vision for behavior. Primates are much more visual animals than rodents and understanding the link between primate visual and medial temporal lobe (MTL) memory structures is an important and largely unexplored open question (*Meister and Buffalo, 2016*). Experimental evidence indicates the existence of functional and anatomical connections between these structures. Functional connections are demonstrated by two principal lines of studies. First, visual behavior is informed by memory as demonstrated by studies of novelty preference in both monkeys and humans (*Wilson and Goldman-Rakic, 1994*; *Manns et al., 2000*; *Jutras and Buffalo, 2010a*). In the novelty preference paradigm, the memory is assessed from looking time: well memorized stimuli are looked at less than novel ones. The specific role of MTL structures in this phenomenon is derived from results showing a decreased novelty preference after MTL lesions or in patients suffering from mild cognitive impairment or Alzheimer’s disease, mental disorders associated with MTL dysfunction (*McKee and Squire, 1993*; *Crutcher et al., 2009*; *Zola et al., 2013*). In monkeys, restricted lesions of hippocampal and/or parahippocampal cortices also decreased novelty preference (*Zola et al., 2000*; *Pascalis et al., 2009*; *Bachevalier et al., 2015*). Second, the link between visual and MTL structures is manifested in coherent neural activities in the two structures. For example, activity of single MTL neurons is modulated by visual saccades (*Sobotka et al., 1997*), the onset of visual stimuli strongly affects hippocampal neural responses (*Jutras and Buffalo, 2010a*) and hippocampal theta oscillations are reset by eye movements (*Jutras and Buffalo, 2010b*; *Hoffman et al., 2013*).

Anatomical connections between visual and memory structures have recently been characterized in the framework of the occipital–parietal–MTL pathway of visuospatial processing (*Kravitz et al., 2011*). There are three principal stages of information processing in this pathway (*Figure 1*A). First, the occipito-parietal circuit processes visual information through visual areas V1-V6 in an egocentric (retinal) frame of reference. Successive information processing in these areas is thought to extract visual features of increasing complexity, including motion and depth cues and relay this information to the parietal cortex. Second, a complex network of interconnected parietal structures relays highly-processed visual cues to support executive, motor and spatial-navigation functions. These structures include the medial, ventral and lateral intraparietal areas (MIP, VIP, LIP) strongly linked with eye movements processing; the middle temporal and medial superior temporal (MT, MST) thought to extract high-level visual motion cues; and the caudal part of the inferior parietal lobule (cIPL), the main relay stage on the way to the medial temporal lobe. The cIPL sends direct projections to the CA1 of the hippocampus as well as to the nearby parahippocampal cortex (PHC). In addition, it sends indirect projections to the same structures via the posterior cingulate cortex (PCC) and the retrosplenial cortex (RSC). Within this complex network, neurons at different neurobiological sites have been reported to code space in a world- or object-centred reference frames (*Galletti et al., 1993*; *Duhamel et al., 1997*; *Snyder et al., 1998*; *Chafee et al., 2007*). Moreover, both PCC and RSC have been repeatedly linked to coordinate transformations between egocentric and allocentric frames of reference (*Vogt et al., 1992*; *Burgess, 2008*; *Epstein and Vass, 2014*). Importantly, information processing in this pathway is strongly affected by directional information thought to be provided by a network of head-direction cells residing in several brain areas, including RSC (*Taube, 2007*). Finally, MTL, and in particular the hippocampus, play a key role in constructing allocentric representations of space in primates (*Hori et al., 2003*; *Ekstrom et al., 2003*).

**Figure 1.**
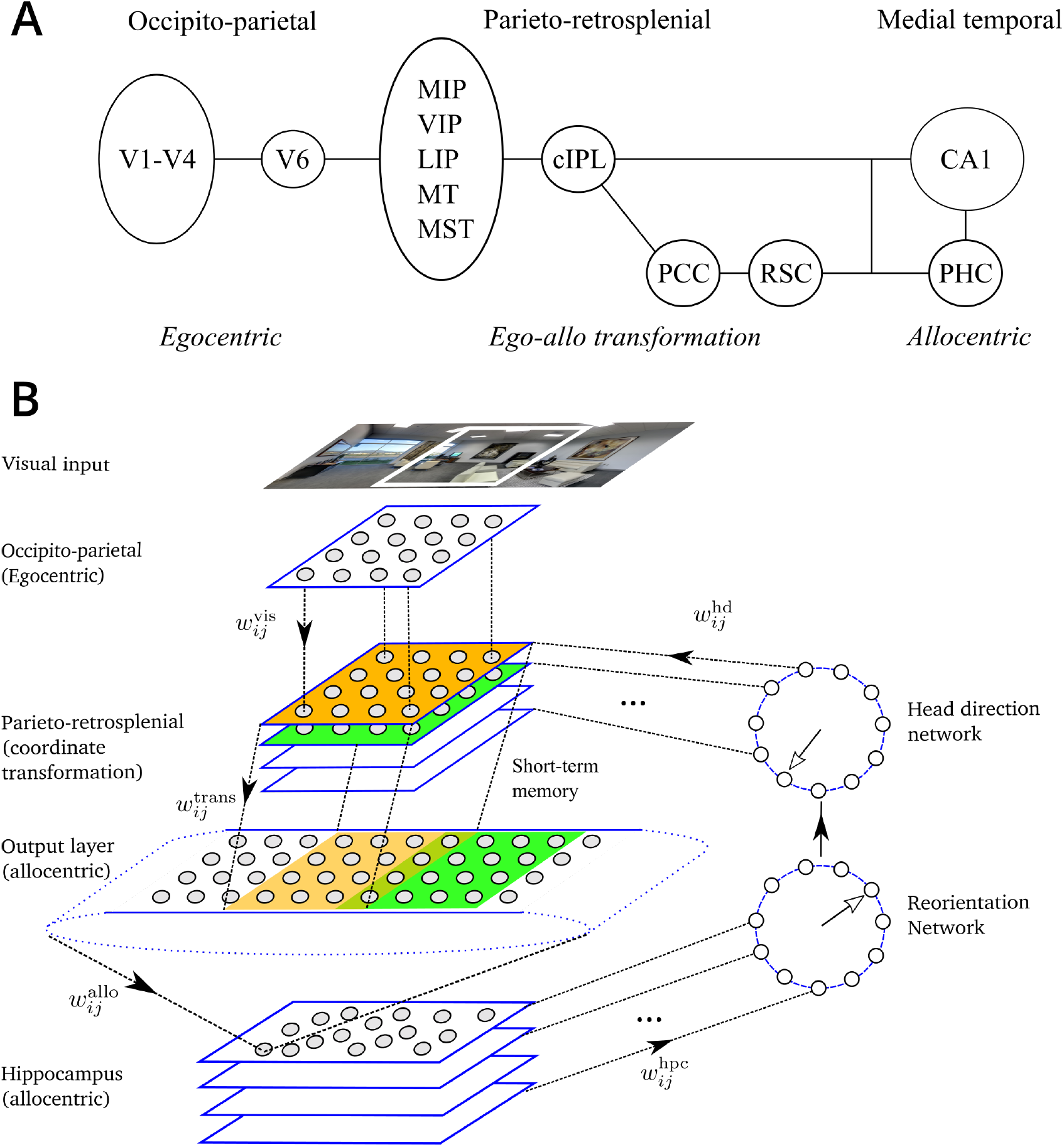
Anatomy and model of the dorsal visual pathway - MTL network. A. Neural structures along the dorsal visual pathway of visuospatial information processing in primates (see text and *Kravitz et al., 2011*). B. Schematic representation of the model. Top image: visual features present in the limited visual field (white square) constitute the visual input of the model. The model network is composed of 6 modules: (1) Occipito-parietal (egocentric); (2) Head-direction network; (3) Parieto-retrosplenial coordinate transformation network; (4) Transformation output layer, which encodes visual features in an allocentric directional frame and spans the full panoramic visual field; (5) Hippocampus; (6) Reorientation network. Projections from the occipito-parietal (visual) areas to the transformation network are topographic. Each head-direction cell activates the corresponding layer of the transformation network. Projections from the different layers of the transformation network to the parieto-retrosplenial output layer are also organized according to head direction: any two layers project topographically to overlapping portions of the output population shifted according to head direction. Synapses between the transformation network and the parietal output network are endowed with short-term memory. Different hippocampal subpopulations project to different neurons in the reorientation network, which in turn corrects head direction signal. Full arrows represent the flow of information in the network. Open arrows represent direction signals in the head direction and reorientation networks.

Given the functional and anatomical connections between visual and memory structures, the question arises as to the nature of neural representations in the dorsal visual pathway. In addition to the well-established role of parieto-retrosplenial networks in coordinate transformations (*Andersen et al., 1993*; *Snyder et al., 1998*; *Salinas and Abbott, 2001*; *Pouget et al., 2002*; *Byrne et al., 2007*), a largely unexplored question concerns the existence of an extra-retinal neural map of the remembered visual space (*Hayhoe et al., 2003*; *Tatler and Land, 2011*; *Land, 2014*). That the task-related visual retinotopic space is remembered has been suggested by studies showing that when asking to recall a recent visual content, eye movements (on a blank screen) closely reflected spatial relations of remembered images (*Brandt and Stark, 1997*; *Johansson and Johansson, 2014*). Moreover, preventing subjects from making eye movements decreased recall performance (*Johansson and Johansson, 2014*; *Laeng et al., 2014*). That not only the retinal egocentric space is remembered but also extra-retinal map of surrounding space is stored in memory is demonstrated in behavioral studies showing that during natural behavior human subjects direct saccades toward extra-retinal locations (*Land et al., 1999*; *Hayhoe et al., 2003*) as well as in fMRI studies suggesting the existence of panoramic visual memory representations (*Park and Chun, 2009*; *Robertson et al., 2016*). Even though suggested by the above studies, the nature of such an extra-retinal map and neural mechanisms underlying its construction and storage in the context of visuospatial orientation are currently unknown.

The present modeling study addresses the question of how such an allocentric representation of surrounding visual space can be constructed and stored by the dorsal visual pathway – MTL networks. We propose that the existence of such a representation relies on short-term memory linking successive egocentric views and we study how the long-term memory of allocentric visual space can affect behavior in spatial and non-spatial experimental paradigms. In particular, our results suggest that allocentric memory effects during spatial reorientation and memory-based visual guidance tasks can be explained by the existence of such a network.

## Methods

The model is a spiking neuron network constructed to mimic information processing steps thought to be performed by the networks along the primate dorsal visual stream, as described above (*Figure 1*A). The model is composed of 6 main modules, or subnetworks (*Figure 1*B). First, the module representing information processing in the occipito-parietal circuit essentially applies a set of Gabor-like orientation filters to the incoming visual images, a standard assumption for basic V1 processing. We do not model eye movements, and assume that a retinotopic visual representation obtained at the level of V1 has been remapped, by the time it arrives to the parietal cortex, to a head-fixed representation by taking into account eye position signals (*Galletti et al., 1993*; *Duhamel et al., 1997*; *Snyder et al., 1998*; *Pouget et al., 2002*). Even though gaze independent, this head-fixed representation is egocentric, or view-dependent, in the sense that it depends on the position and orientation the modeled animal (i.e., its head) in space. Second, we model the directional sense by a ring of cells whose activity is approximately Gaussian around their preferred orientations (*Taube, 2007*) and that is sending projections to the parietal cortex (*Brotchie et al., 1995*; *Snyder et al., 1998*). Third, both the activities of the egocentric network and the head direction signal converge onto the network modeling the role of the parieto-retrosplenial network in coordinate transformation. This transformation network uses head direction to convert egocentric view-dependent representations to a head-orientation-independent, or world-fixed one. The coordinate transformation is performed essentially by the same mechanism as the retinotopic-to-head-fixed conversion mentioned above, but in contrast to previous models it does so using low-level topographically-organized visual information. Fourth, the resulting head orientation-independent visual representation can be referred to as spatiotopic, or allocentric, since its constituent visual features are coded in a world-fixed directional reference frame. The existence of such a representation is one of the central claims of the present work and it is proposed to serve as a neural interface between sensory and mnemonic structures along the medial-temporal division of the dorsal path network. Fifth, the allocentric output of the parieto-retrosplenial network arrives to the hippocampus, modeled by a network of cells that learn, by a competitive mechanism, allocentric visual patterns provided by the parietal network. As will be clear from the following, in the context of spatial navigation these cells can be considered as place cells, whereas in a non-spatial context they can be considered as representing memorized visual stimuli. Finally, the reorientation module associates allocentric memories with an associated directional reference frame and feeds back to the head direction cells. The activity of this network represents the correction signal for self-orientation. When the memorized information corresponds to the newly arrived one, the correction signal is zero, whereas in the case of disorientation or in response to specific manipulations of visual cues, it can provide fast adjustment of the self-orientation signal. In the Results section we show that a similar reorientation mechanism can be responsible for behavioral decisions in spatial, as well as non-spatial tasks in primates.

### Occipito-parietal input circuit

The occipito-parietal network is modeled by a single rectangular sheet of *N*_x_ × *N*_y_ visual neurons, uniformly covering the visual field. In all simulations, except Simulation 6 below, the size of the visual field was limited to 160 × 100°, approximately representing that of a primate. The activities of these visual neurons are computed in four steps. First, input images are convolved (using OpenCV `filter2D()` function) with Gabor filters of 4 different orientations (0, 90°, 180°, 270°) at 2 spatial frequencies (0.5 cpd, 2.5 cpd), chosen so as to detect visual features in simulated experiments. Second, the 8 convolution images are discretized with *N*_x_ × *N*_y_ grid, and the maximal response at each position is chosen, producing an array of *N*_x_*N*_y_ filter responses. These operations are assumed to roughly mimic retinotopic V1 processing (*Heeger, 1992*), transformed into a head-fixed reference frame using eye-position information. Third, the vector of filter activities at time *t* is normalized to have maximal value of unity. Fourth, a population of *N*_vis_ = *N*_x_*N*_y_ Poisson neurons is created with mean rates given by the activity of the corresponding filters scaled by the constant maximal rate *A*_vis_ (see ***Table 1*** for the values of all parameters in the model). For a Poisson neuron with rate *r*, the probability of emitting a spike during a small period of time *δt* is equal to *rδt* (*Gerstner et al., 2014*).

### Head direction

The head direction network is composed of *N*_hd_ = 36 Poisson neurons organized in a circle, such that neurons’ preferred directions *φ*_*k*_ are uniformly distributed between 0 and 2*n*. The tuning curves of the modeled head-direction neurons are Gaussian with maximum rate *A*_hd_ and width (*J*_hd_ = 8°. Thus, the rate of head-direction neuron *k* when the model animal’s head is oriented in the direction *φ* is given by

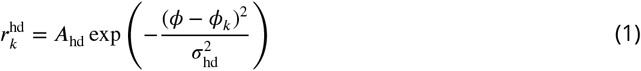

Such a network generates a Gaussian activity profile centered around *φ*. Our model does not explicitly implement a line attractor dynamics hypothesized to support head direction signal (*Zhang, 1996*), but it is consistent with it. Head direction cells have been found in several brain areas in rodents and primates (see *Taube, 2007*, for review), and there is evidence that parietal cortex receives head direction signals (*Brotchie et al., 1995*).

### Parietal transformation network

The parietal transformation network is inspired by previous models (*Becker and Burgess, 2001*; *Byrne et al., 2007*) but in contrast to them it operates directly on activities of the Gabor-like visual cells. The transformation of coordinates between the head-fixed and world-fixed coordinates is performed by multiple subpopulations of leaky integrate-and-fire (LIF) neurons organized as twodimensional layers of neurons (see *Figure 1*). Neurons in each layer of the transformation network are in a one-to-one relationship with the visual population and so at each moment *t* each transformation layer receives a copy of the egocentric (head-fixed) visual input. Therefore, the number of neurons in each transformation layer is equal to *N*_*vis*_. Apart from the visual input, the transformation network also receives input from the population of head direction cells. There is a topographic relationship between the sub-populations of the transformation network and different head directions: each head-direction cell sends excitatory projections to neurons only in one associated layer of the transformation network. Thus, an input from head-direction cells strongly activates only a small subset of transformation layers which transmit visual information to the downstream population. More specifically, only the layers which are associated with head directions close to the actual orientation of the head are active. The number of layers in the transformation network is then equal to *N*_hd_, giving the total number of neurons in the transformation network *N*_trans_ = *N*_vis_*N*_hd_.

Thus, in a *k*-th layer of the transformation network, the membrane potential *v*_*i*_(*t*) of the -th LIF neuron is governed by the following equation (omitting the layer index for clarity):

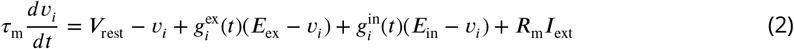

with the membrane time constant *τ*_*m*_, resting potential *V*_rest_, excitatory and inhibitory reversal potentials *E*_ex_ and *E*_in_, as well as the membrane resistance *R*_m_. When the membrane potential reaches threshold *V*_th_, the neuron fires an action potential. At the same time, *v* is reset to *V*_reset_ and the neuron enters the absolute refractory period Δ_abs_ during which it cannot emit spikes. A constant external current *I*_ext_ is added to each neuron to simulate baseline activity induced by other (unspecified) neurons from the network.

The excitatory conductance in transformation network neurons depends only on the visual input (and thus is independent from index *k*). It is modeled as a combination of *α*-amino-3-hydroxy-5-methyl-4-isoxazolepropionic acid (AMPA) and N-methyl-d-aspartate (NMDA) receptors activation 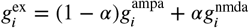, that are described by (*Murray et al., 2014*):

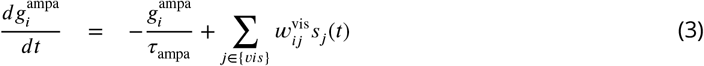

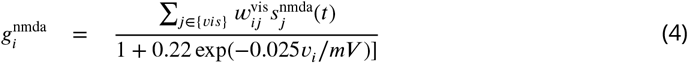

where the index *j* runs over input (visual) neurons connected to it, 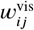 are the connection weights and *s*_*j*_(*t*) = 1 if a presynaptic spike arrives at time *t* and *s*_*j*_(*t*) = 0 otherwise. Constant *r*_ampa_ determines the time scale of AMPA receptor activation. The NMDA-receptor-mediated current exhibits voltage dependence controlled by magnesium concentration and the NMDA gating variable 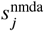 evolves according to the following equations:

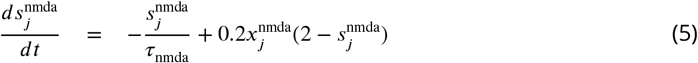

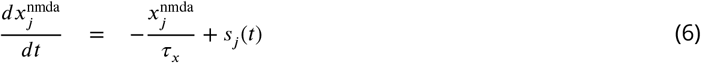

where *τ*_nmda_ and *r*_*x*_ re the decay time constants (*Murray et al., 2014*).

The inhibitory conductance in these neurons depends only on the head-direction cells and ensures that a small subset of transformation layers (i.e. those associated with nearby head directions) are active. To implement it, we employ a simple scheme in which all transformation layer neurons are self-inhibitory, and this inhibition is counteracted by the excitatory input from the head-direction cells. Thus, the inhibitory conductance of the -th neuron in the *k*-th layer is given by

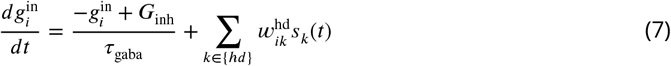

where *G*_inh_ is the constant maximum amount of self-inhibition and 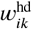 are the synaptic weights of connections from the head-direction cells. In the current implementation, there is one-to-one correspondence between the head-direction cells and the layers of the transformation network, so *w*_*ik*_ = 1 only for an associated head-direction cell *φ*_*k*_ and 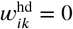 otherwise.

#### Learning the weights in the transformation network

The connection weights 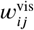 from the visual neurons to the parietal transformation cells and 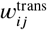 from the parietal transformation cells to the parietal output neurons are assumed to be learned during development by a supervised mechanism, similar to the one proposed to occur during sensory-motor transformation learning (*Zipser and Andersen, 1988*; *Salinas and Abbott, 1995*). In this models it is proposed that when an object is seen (i.e. its retinal position and an associated gaze direction are given), grasping the object by hand (that operates w.r.t. the body-fixed reference frame) provides a teaching signal to learn the coordinate transformation. A similar process is assumed to occur here, but instead of learning body-based coordinates using gaze direction, the model learns world-fixed coordinates using head direction.

More specifically, synaptic weights in the coordinate-transformation network were set by the following procedure. First, the network was presented with an edge-like stimulus at a random orientation and at a randomly chosen location in the visual field. Second, upon the stimulus presentation, the head direction was fixed at a randomly chosen angle *φ*. Third, neurons in the transformation layers associated with the chosen head direction were activated with the average firing rates equal to the rates of the corresponding visual neurons, while neurons in the parietal output layer were activated with the same average rates but shifted according to the chosen head direction (representing the teaching signal). Fourth, the synaptic weights in the network were set according to the Hebbian prescription:

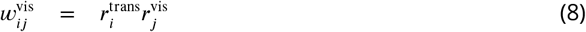

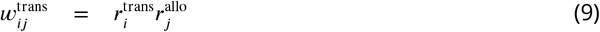

where 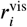, 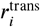 and 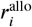 are the mean firing rates of the corresponding visual neurons, transformation network neurons and parietal output neurons, respectively. Fifth, the weight vector of each neuron was normalized to have the unity norm. This procedure has been performed for edge-like stimuli at 4 different orientations (corresponding to 4 Gabor filter orientations), placed in the locations spanning the whole visual field and at head directions spanning 360°. Synaptic weights (*Equation 8*-***9***) were fixed to the learned values prior to all the simulation presented here. No updates were performed on these weights during the simulations.

### Parietal output population

All layers of the transformation network project to the parietal output population, which codes image features in an allocentric (world-fixed) directional frame. The parietal output population is represented by a two-dimensional neuronal sheet spanning 360 × 100°, i.e. a full panoramic view. It is encoded by a grid of 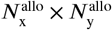 neurons. Each layer of the upstream transformation network projects to a portion of the sheet according to the head direction associated with this layer (see *Figure 1*). Since any two nearby layers of the transformation network are associated with head directions shifted relative to each other by 360°/*N*_hd_ = 10°, the overlap between their projections on the parietal output layer is 140° (given that the size of the limited visual field is 160°).

At each moment in time, a spiking representation of the current visual stream (i.e. a spiking copy of the visual input, gated by the head direction cells) arrives to the allocentric neurons spatially shifted according to the current head direction. For example, if two egocentric views (each spanning 160°) are observed at head directions −45° and 45° with respect to an arbitrary north direction, these two views arrive at the allocentric population spatially shifted relative to one another by 90°, so that the activated neurons in the allocentric population span 230°. To ensure that subsequent snapshots are accumulated in time (e.g. during head rotation), the synapses between neurons in the transformation layers and the allocentric population are endowed with short-term memory, implemented by a prolonged activation of NMDA receptors (*Durstewitz et al., 2000*). Such synapses result in a sustained activity of allocentric output neurons during a period of time sufficient for downstream plasticity mechanism to store information from accumulated snapshots.

Thus, the membrane potential of the -th neuron in the allocentric output population is governed by *Equation 2* with the synaptic conductance terms determined as follows. First, the excitatory AMPA conductance is given by *Equation 3* but with the input provided by transformation network neurons via weights 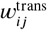. Second, the NMDA conductance is described by *Equation 4*, but with the synaptic time scale increased by a factor of 6. This is done to ensure sustained activation of the output neurons upon changes in the visual input. Third, inhibitory input is set to zero for these neurons.

### Hippocampal neurons

As a result of the upstream processing, neuronal input to the hippocampus represents visual features in an allocentric directional frame. Neurons in the parietal output population are connected in an all-to-all fashion to the population of modeled hippocampal cells and the connection weights that are updated during learning according to an spike-timing-dependent plasticity (STDP) rule below. In addition, lateral inhibition between hippocampal neurons ensures a soft winner-take-all dynamics, such that sufficiently different patterns in the visual input become associated with small distinct subpopulations of hippocampal neurons.

Thus, the membrane equation of the -th hippocampal neuron is given by *Equation 2*. The excitatory conductances are given by *Equation 3*-***4***, but with the input provided by the parietal output neurons via weights 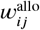. Upon the initial entry to a novel environment these weights are initialized to small random values. During learning, the amount of synaptic modification induced by a single pair of pre- and post-synaptic spikes is given by

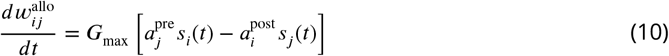

where *s*_*i*_ (*t*) and *s*_*j*_(*t*) detect pre- and post-synaptic spikes, respectively, and

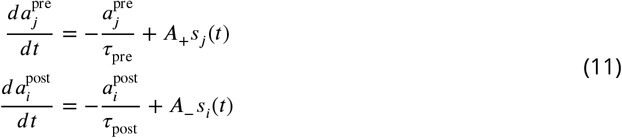

The inhibitory conductance of the *i*-th neuron is governed by the following equation:

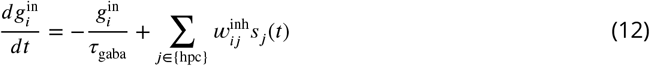

in which *τ*_gaba_ determines the time scale of synaptic inhibition as before, and weights 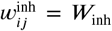 are constant and ensure that each hippocampal neuron inhibits all other hippocampal neurons proportionally to its activity.

The real hippocampal circuit is complex and consists of several interconnected populations of cells with various spatially-tuned activity profiles (*Moser et al., 2017*). In the present simple model of hippocampal activity we consider only CA1 – the first stage of hippocampal processing where sensory cues give rise to spatially selective firing and which receives direct projections from the entorhinal cortex, an input gateway to the hippocampus. See our recent work (*Li et al., 2020*) for a computational model of how such a vision-based representation can be further integrated with self-motion cues to build neural representations of complex environments.

### Reorientation network

During one continuous experimental trial (e.g. an exploration trial in a novel environment or an observation of a novel image on the screen), the reference frame for head direction is fixed and all processing operations in the network are performed with respect to the origin of this reference frame. An allocentric information stored by the hippocampus as a result of the trial can be correctly used for future action only if the origin of the reference frame is stored with it. Therefore, if in a subsequent trial the actions to be performed require memory of the previous one, the network should be able to recover the original directional reference (this of course should happen only when the visual information received at the start of the trial is considered familiar). Reorientation is defined here as the process by which the origin of the stored reference frame is recovered.

In our model, reorientation is implemented by a network of neurons organized similarly to the head direction network described earlier. This reorientation network is proposed to reside in the RSC and to be driven by the direct hippocampal input. We model the component of this process that is automatic, fast, bottom-up, and does not require costly object/landmark processing. A large body of reorientation studies in many animal species including primates and human children show that object identities are often ignored during reorientation (*Cheng and Newcombe, 2005*). The conditions in which most of these reorientation studies were performed usually are such that there is no single conspicuous point-like cue in the environment that can be reliably associated with a reference direction. For example, in many studies the directional cues come from the geometric layout of an experimental enclosure. Lesion studies in rats suggest that reorientation in these conditions requires an intact hippocampus (*McGregor et al., 2004*). Furthermore, we propose that this reorientation network is active all the time, in contrast to being consciously “turned on” when the animal “feels disoriented”. Therefore, we expect that its effects can be observed even when no specific disorientation procedure was performed. In particular, we suggest in the Results that a manipulation of objects on the screen can result in automatic corrections of directional sense that can be observed during visual search.

The reorientation network consists of *N*_re_ neurons with ring-like topology whose preferred positions are uniformly distributed on a circle. Therefore, the difference between preferred position of two nearby reorientation cells is Δ*φ* = 2*π*/*N*_re_. The membrane potential of the -th reorientation neuron is described by the LIF equation (*Equation 2*). Excitatory conductances are described by *Equation 3*-***4*** with the input to the neuron provided by hippocampal place cells via weights 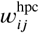. There is no inhibition in the network, and so the inhibitory conductance is set to 0. The ability of the network to perform reorientation is determined by afferent connection weights from the hippocampal cells, which are determined as follows.

Since all visual information observed during a trial is linked to the same directional frame, all hippocampal cells learned during the trial are connected to a single neuron of the reorientation network, the one with the preferred direction 0° (*Figure 2*). The connection weights between the hippocampal cells and the neuron are updated using STDP rule, *Equation 10*-***11*** (this is not essential for the model to work, so that setting the weights to a constant value will give similar results). Once the training trial is finished, *N*_*re*_ copies of the learned hippocampal population are created, each corresponding to a separate neuron in the reorientation network. In each copy, all cells have the same input and output weights as the corresponding cells in the original population, but their connection profile is different. In particular, the copy corresponding to the reorientation neuron with preferred direction Δ*φ* is connected to pre-synaptic cells that are shifted by the same angle in the topographically-organized allocentric layer (*Figure 2*). In machine learning literature, this technique is called “weight sharing” and it allows to achieve translation invariance for object recognition in images. Here, we apply a similar technique in order to detect familiar snapshots and the directional reference frame associated with them.

**Figure 2.**
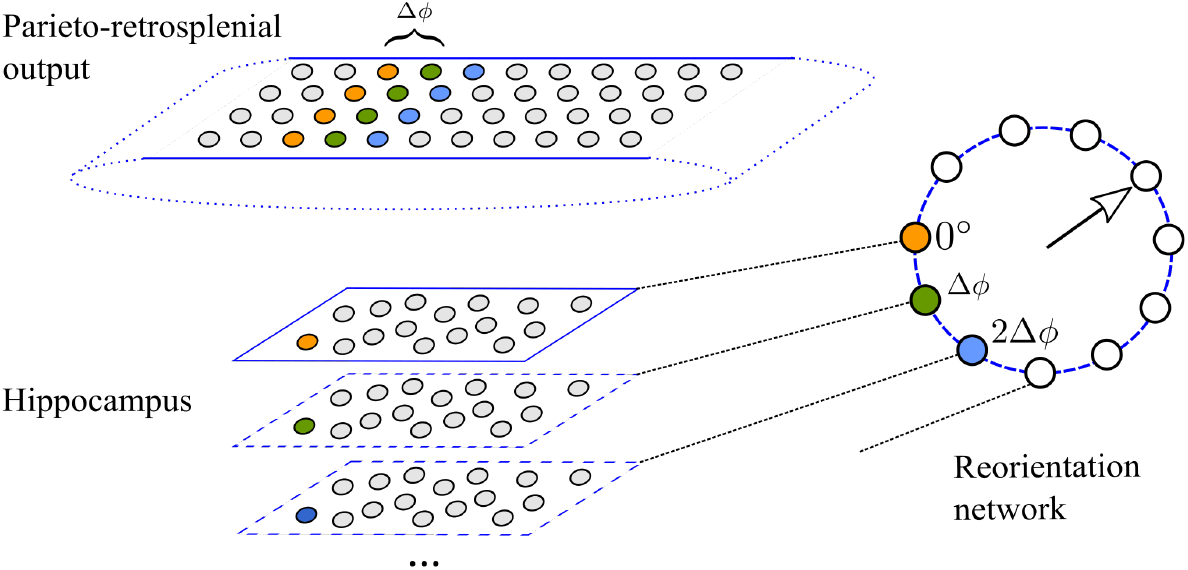
Implementation of the reorientation network. Top: the output population of the parieto-retrosplenial network. Bottom: hippocampal cells. The population outlined by full lines is the original population learned during training. As a result of learning, the hippocampal cell shown in orange is connected to the presynaptic cells of the same color (connection weights not shown). All hippocampal cells in the original population are connected to a single cell (0°) in the reorientation network (Right). The hippocampal populations outlined by the dashed lines are copies of the original population that implement weight sharing: e.g. the hippocampal cell shown in green (blue) has the same connection weight values as the orange cell, but it is connected to pre- and post-synaptic cells shifted by Δ*φ* (2Δ*φ*) with respect to those of the orange cell. The number of copies of the original hippocampal population is thus the same as the number of neurons in the reorientation network.

Suppose, for example, that as a result of learning during a trial, a hippocampal cell is associated with a vertical edge in the environment represented by 4 presynaptic cells in the output layer of the transformation network (cells shown in orange in *Figure 2*). Suppose further that during an inter-trial interval the head direction network has drifted (or was externally manipulated), so that at the start of the new trial the internal sense of direction is off by 2Δ*φ*. When the animal sees the same visual pattern again, it will be projected onto the allocentric layer shifted by the same amount due to the error in orientation (blue cells in *Figure 2*). This will in turn cause the hippocampal subpopulation that includes the blue cell to be most strongly active in response to the visual input, and so the activity peak of the reorientation network will signal the orientation error (2Δ*φ*). The reorientation is then performed by readjusting the head direction network to minimize the orientation error. In the current implementation this is done algorithmically by subtracting the error signal from the actual head direction, but it can also be implemented by attractor dynamics in the head direction layer.

### Simulation details

The spiking artificial neural network model described above was implemented using Python 2.7 and Brian 2 spiking neural network simulator (*Stimberg et al., 2019*). The time step for neuronal simulation was set to 1 ms, while the sampling rate of visual information was 10 Hz, according to the proposals relating oscillatory brain rhythms in the range 6–10 Hz to information sampling (*Hasselmo et al., 2002*; *Busch and VanRullen, 2010*). At the start of each simulation, the weights 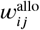 and 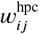 were initialized to small random values (the other weights were trained as described earlier and fixed for all simulations), see *Figure 1*B. Parameters of the model are listed in *Table 1*, and the sections below provide additional details of all simulations.

**Table 1.**
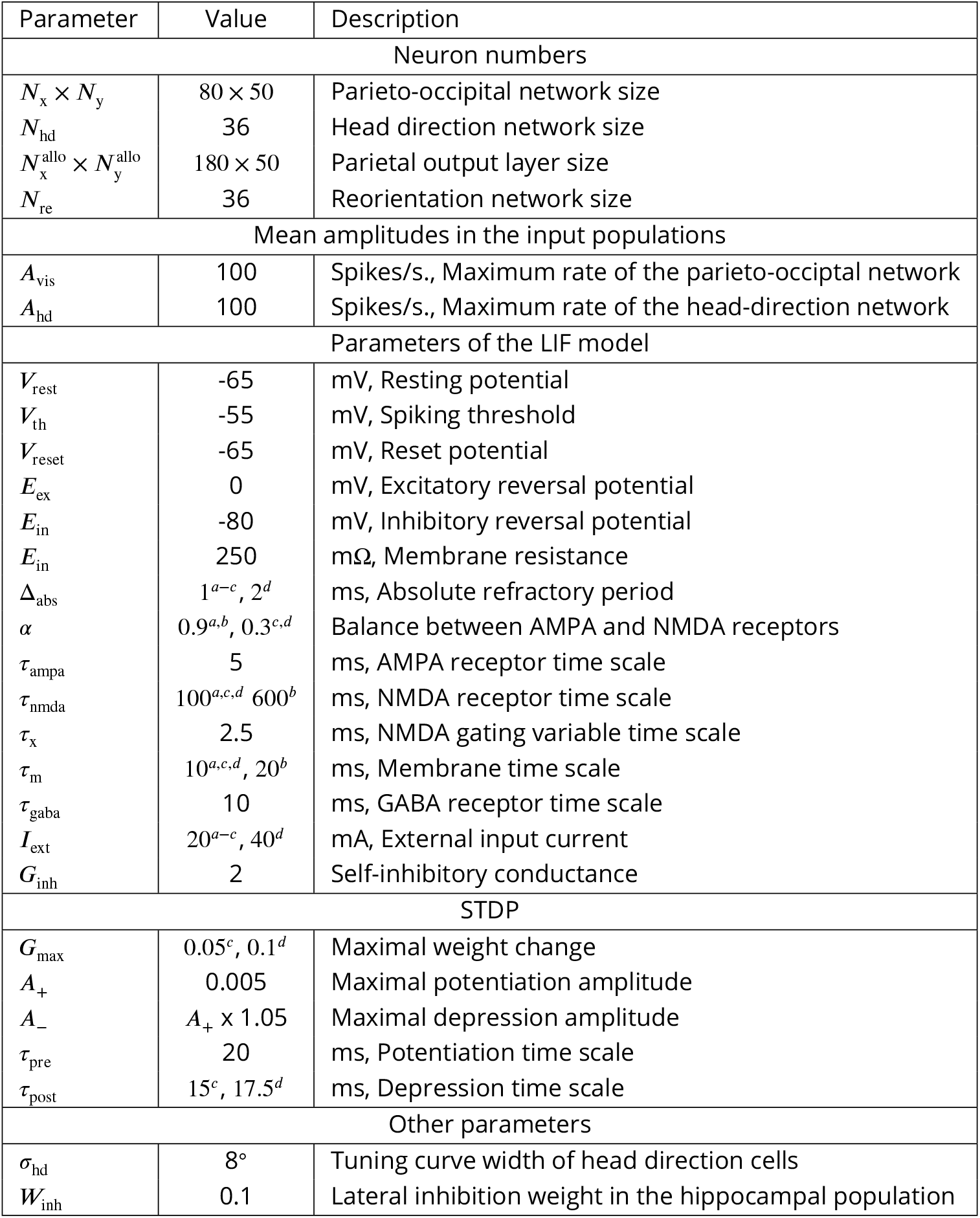
Parameters of the model. a, Occipito-parietal circuit. b, Parieto-retrosplenial transformation network. c, Hippocampus. d, Reorientation network.

#### Simulation 1: Egocentric-allocentric transformation

The first simulation was inspired by the study of ***Snyder et al.*** (***1998***), in which monkeys observed visual stimuli at identical retinal locations, but for different orientations of the head with respect to the world, in order to assess whether parietal neurons were modulated by the allocentric head direction. Thus, in this simulation, the head direction angle *φ* was varied from −50° to 50° in 100 sessions. For each trial of a session, the mean rates of the head-direction neurons were calculated according to *Equation 1* and fixed for the rest of the trial. The stimulus (vertical black bar, width: 10°) was shifted horizontally along the midline of the visual field (160 × 100°) from left to right in 1° steps, such that it remained at each position for 100ms. The neuronal spikes were recorded from the occipito-parietal network, the parieto-retrosplenial transformation network and its output layer, for each stimulus position across 10 trials per session. Mean firing rates were then calculated from these data.

#### Simulation 2: Accumulation of successive views using short-term synaptic memory

The aim of the second simulation was to illustrate the synaptic mechanism for an integration of successive visual snapshots in time, instrumental for spatial coding. We model a monkey that remains in the same spatial location and turns its head from left to right. Thus, the model was presented with a set of 9 successive overlapping views (160 × 100°) taken from a panoramic (360 × 100°) image, 100ms per view. Initial head direction was arbitrarily set to 0°.

#### Simulation 3: Encoding of allocentric visual information during spatial exploration

In the third simulation we studied the role of temporal accumulation of visual information for spatial coding. The model ran through a square 3D environment (area: 10×10 m, wall height 6 m) for about 10 min so as to cover uniformly its area. The visual input was provided by a cylindrical camera (160 × 100°) placed at the location of the model animal and oriented according to its head direction. At each spatial location, 9 successive views of the environment were taken in different directions (as in the Simulation 2). The vector of mean firing rates of the occipito-parietal neurons at a single spatial location and orientation constituted the egocentric population vector. The mean firing rates of the the parieto-retrosplenial output neurons at each location constituted the allocentric population vector (this population vector is independent from orientation as a result of coordinate transformation). To compare spatial information content in the two populations, we first estimated intrinsic dimensionality of the two sets of population vectors. This was performed using two recent state-of-the art methods: DANCo (*Ceruti et al., 2014*), as implemented by the `intrinsicDimension` R package, and ID_fit (*Granata and Carnevale, 2016*). For both methods, the principal parameter affecting dimensionality estimation is the number of neighbors for each point in the set that is used to make local estimates of the manifold dimension. Second, we used two different methods to visualize the structure of the low-dimensional manifold: Isomap (*Tenenbaum et al., 2000*) and t-SNE (*van der Maaten and Hinton, 2008*). To extract principal axes of the manifold, we used PCA on the data points projected on two principal dimensions provided by the above methods. We chose the parameter values for which the visualized manifold best approximates the original space. We then determined a set of points (i.e. population vectors) that lie close to the principal axes of the manifold and visualized them in the original environment. If the manifold structure corresponds well to the spatial structure of the underlying environment, the principal axes of the manifold should lie close to the principal axes of the environment.

#### Simulation 4: Visual responses of hippocampal neurons in an image memorization task

This simulation was inspired by the study of *Jutras and Buffalo* (***2010a***) in which a large set of novel visual stimuli was presented to monkeys on a computer screen. Neuronal activity in the hippocampal formation in response to the visual stimuli was recorded. One of the results of this study suggested that hippocampal neurons encode stimulus novelty in their firing rates. To simulate this result, we presented to the model 100 novel stimuli randomly chosen from the dataset retrieved from http://www.vision.caltech.edu/Image_Datasets/Caltech101). The stimuli (resized to 160 × 100 pixels) were shown to the model successively in one continuous session (500ms stimulus presentation time + 1000ms inter-trial interval with no stimuli) and the activities of the hippocampal neurons during learning were recorded.

#### Simulation 5: Spatial reorientation

In this simulation of the experiment of ***Gouteux et al.*** (***2001***), the testing room was a rectangular 3D environment with area 20×10 m and wall height 6m. In the “No cues” task the only visual features in the room were provided by the outlines of the walls. In the other 3 tasks, a square visual cue was presented in the middle of one of the walls with the edge length equal to 1/6 (small cue), 1/3 (medium cue) or 1/2 (large cue) of the environment width. Each task consisted of two phases, exploration and reorientation. During the exploration phase the modeled animal uniformly explored the environment, as in Simulation 3. The reorientation phase composed multiple trials. At the beginning of each trial, the model was placed at one of spatial locations covering the environment in a uniform grid. At each of these locations, 9 successive views were taken. Reorientation performance was assessed in two ways: (i) only the first view at each location was used for reorientation; (ii) successive views accumulated over 60 successive positions were used for reorientation.

#### Simulation 6: Memory-based visual search

In this simulation we used a dataset of visual images used in the study by ***Fiehler et al.*** (***2014***). This dataset consists of 18 image sets corresponding to 18 different arrangements of the same 6 objects (mug, plate, egg, jam, butter, espresso cooker). Each set includes a control image (all objects on the table in their initial positions) and images in which one of the objects is missing (target object) and one or more other objects displaced to the left or to the right. In the simulation we used only a subset of all images in a set that included either 1, 3 or 5 of the objects mentioned above displaced either to the left or to the right (referred to as “local” condition in *Fiehler et al., 2014*), giving rise to 6 experimental conditions. In each condition, there were 18 test images of displaced objects, plus the associated control images. Taking into account the distance between the animal and the screen as well as the size of the image (provided in *Fiehler et al., 2014*), we calculated the size of the image in degrees of visual field. We then determined a rectangular portion of the image (30 × 15°) that included all objects in initial and displaced positions in all images. The contents of this area served as an input to the model. Thus, in this simulation the spatial resolution of the visual input was higher than in the previous simulations as the visual field of the model was smaller, but the size of the input network was kept the same.

During each simulation trial, the image of objects in initial positions was first presented to the network during 2000 ms and stored by the hippocampal cells. The image of displaced objects (in one of the 6 conditions above) was subsequently presented to the network for the same amount of time and the orientation error was read out from the mean firing rates of the reorientation network.

## Results

### Input and output parietal model neurons encode sensory representations in distinct reference frames

We start with a characterization of modeled dorsal visual stream neurons in the case when a simulated animal is assumed to sit in front of a screen and is free to rotate its head (*Duhamel et al., 1997*; *Snyder et al., 1998*, for simplicity, we assume that rotation occurs only in the horizontal plane). The firing rate of occipito-parietal (input) neurons and the output parietal neurons as a function of the allocentric position of a visual stimulus (i.e. a vertical bar moving horizontally across the visual field, see Methods) was measured for two different head directions (*Figure 3*A,B). For a neuron in the input population, a change in head direction induces the corresponding change of the receptive field of the neuron, since its receptive field shifts together with the head along the allocentric position axis (*Figure 3*C). In contrast, for a parietal output neuron, a change in head direction does not influence the position of its receptive field, which remains fixed in an allocentric frame (*Figure 3*D). To show that this is also true on the population level, we measured, for all visual input cells and all parietal output cells, the amount of shift in its receptive field position as a function of head direction shift, while the head was rotated from −50° to 50°. For cells in the occipito-parietal visual area, the average linear slope of the dependence is close to 1, whereas in the allocentric parietal population the average slope is close to 0 (*Figure 3*E), meaning that these two populations encode the visual stimulus in the two different reference frames: head-fixed and world-fixed. These properties of model neurons reproduce well-known monkey data showing that different sub-populations of parietal cortex neurons encode visual features in the two reference frames (*Duhamel et al., 1997*; *Snyder et al., 1998*).

**Figure 3.**
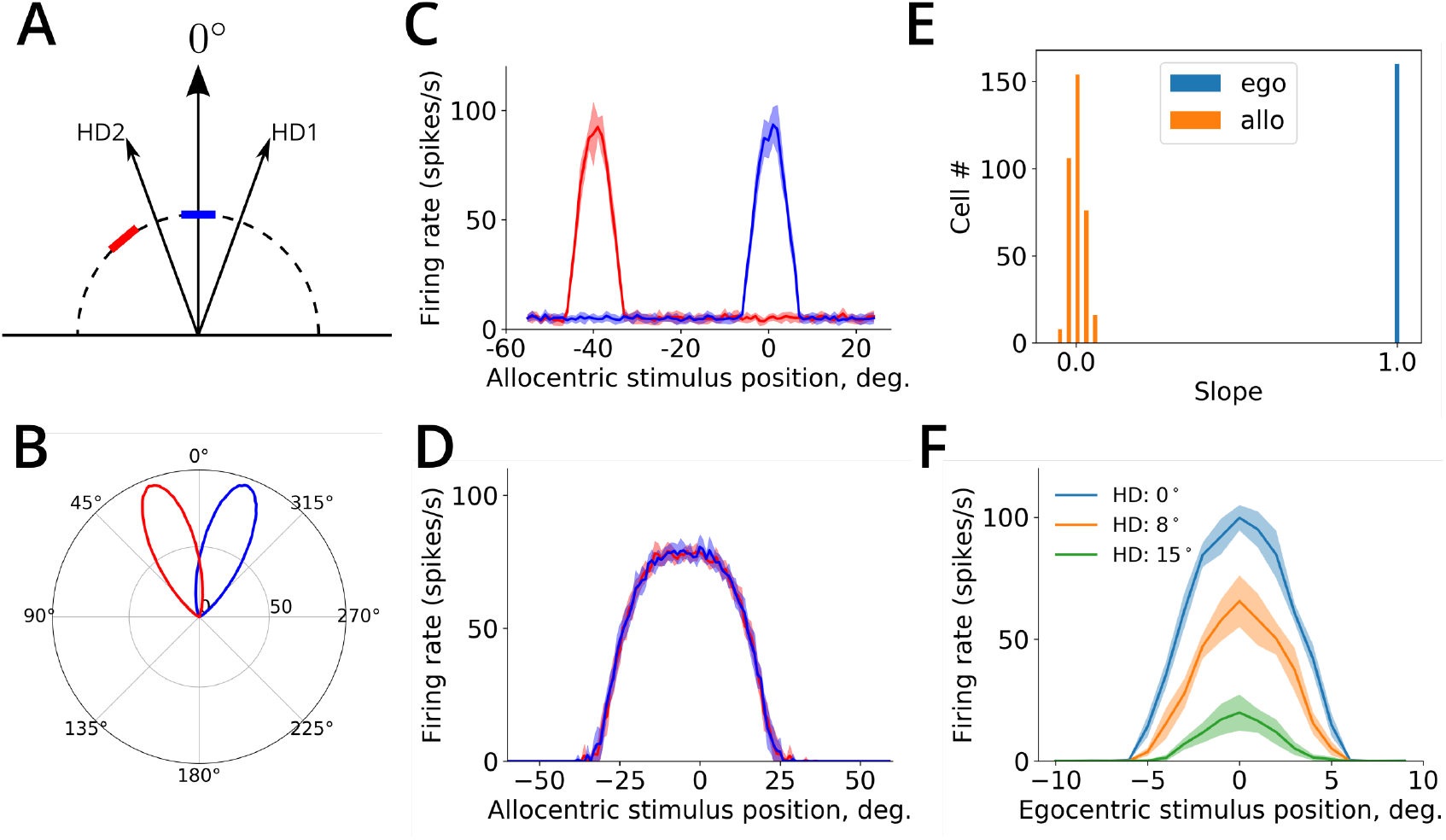
Properties of neurons in the coordinate-transformation network. A. A schematic representation of the receptive field of one input visual input neuron at two head directions (HD1 and HD2). The position of the receptive field of the neuron is shown by the blue and red bar for HD1 and HD2, respectively. B. The population activity of head direction cells in the model at 20° (HD1) and −20° (HD2). C. Tuning curves of an input visual neuron (±SD) as a function of the allocentric stimulus position for the two head directions represented in B. D. Tuning curves of an allocentric output neuron for the same two head directions. E. Histograms show the distributions of the linear dependence slopes between the shift in the receptive field position and the shift in head direction, for egocentric (in blue) and allocentric (in orange) neuronal populations. F. Tuning curves of the same parietal transformation neuron as a function of the egocentric stimulus position for three different head directions are shown. Head direction information is encoded in the activity gain the neuron.

The receptive fields of the intermediate neurons of the coordinate transformation network exhibit gain modulation by head direction (*Figure 3*F), as do monkey parietal neurons (*Snyder et al., 1998*). The hypothesis of reference-frame conversion via gain modulation has been extensively studied in both experimental and theoretical work, in the context of sensory-motor coordination during vision-guided reaching (*Pouget and Sejnowski, 1997*; *Salinas and Abbott, 2001*; *Avillac et al., 2005*). While coordinate-transformation processes involved in the two cases are conceptually similar, the underlying neuronal computations can differ substantially, because the former requires simultaneous remapping for the whole visual field, while the latter is limited to the computation of coordinates for a single target location (i.e. a representation of the point-like reaching target). This difference limits the use of noise-reducing attractor-like dynamics that is an essential component in point-based sensory-motor transformation models (*Pouget et al., 2002*), because in full-field transformation the information and noise are mixed together in a single visual input stream.

### Spatial coding using temporal accumulation of successive views

Because of a limited view field, at each moment in time the simulated animal can directly observe only a restricted portion of the surrounding visual environment (i.e. a visual snapshot, see *Figure 4*A,B). That these snapshot-like representations are represented in memory, has been demonstrated in a number of studies showing viewpoint-dependent memory representations (*Diwadkar and McNamara, 1997*; *Christou and Bülthoff, 1999*; *Gaunet et al., 2001*). Moreover, experimental evidence suggests that visual information can be accumulated from successive snapshots during e.g. head rotation, giving rise to a panoramic-like representation of the surrounding environment that can inform future goal-oriented behavior (*Tatler et al., 2003*; *Oliva et al., 2004*; *Park and Chun, 2009*; *Robertson et al., 2016*). A candidate neural mechanism for implementing such integration is short-term memory, i.e. the ability of a neuron to sustain stimulus-related activity for a short period of time (*Goldman-Rakic, 1995*). In our model, this is implemented by sustained firing via prolonged NMDA receptor activation (*Figure 4*C,D). Combined with STDP learning rule in the connections between the parietal output neurons and the hippocampus, this mechanism ensures that a time-integrated sequence of visual snapshots is stored in the synapses to hippocampal neurons. In particular, head rotation results in a temporarily activated panoramic representation in the population of output parietal neurons that project to CA1. STDP in these synapses ensures that these panoramic representations are stored in the synapses to downstream hippocampal neurons (*Figure 4*E).

**Figure 4.**
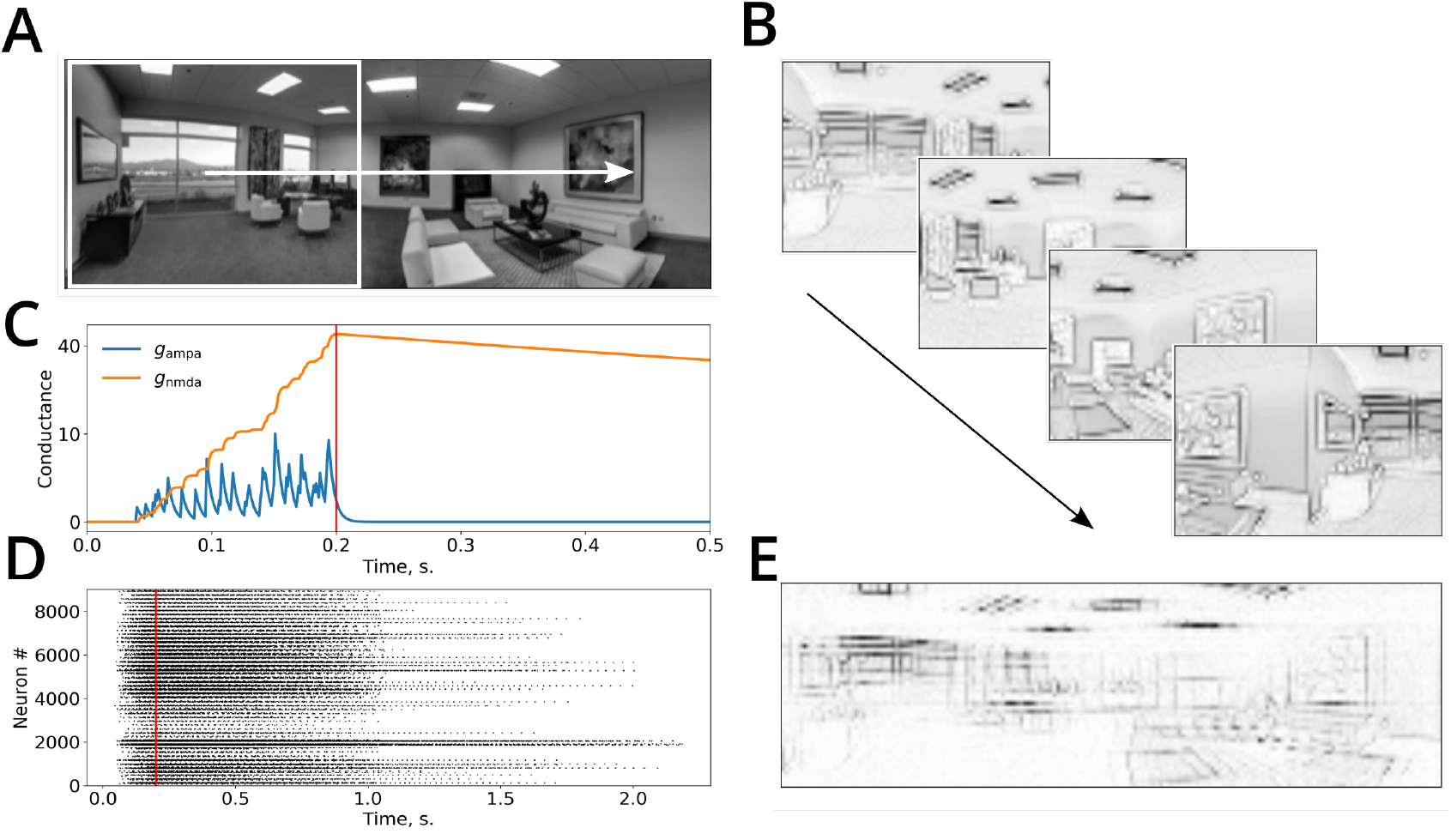
Temporal accumulation of successive visual snapshots in the model. A. A panoramic image of an environment superimposed with the visual field of the simulated animal (white rectangle). The white arrow shows the direction of visual scan path. B. Several successive visual snapshots along the scan path shown in A are represented by mean firing rates of the occipito-parietal (egocentric) network. C. An example of the evolution of AMPA and NMDA receptor conductances of parieto-retrosplenial output neurons as a function of time. Stimulus onset: *t* = 0, stimulus offset: *t* = 200 ms (red line). D. Raster plot of spiking activities of the output neurons showing short-term memory in this network. An input is presented at time 0 and is switched off at the time shown by the red vertical line. The neurons remain active after stimulus offset due NMDA-receptor mediated short-term memory. E. As a result of learning, synaptic weight matrix of a single hippocampal neuron stores the activity of the parieto-retrosplenial output layer accumulated over several successive snapshots shown in B.

A large amount of experimental evidence suggests that many animal species encode a geometric layout of the surrounding space (*Cheng and Newcombe, 2005*; *O’Keefe and Burgess, 1996*; *Gouteux et al., 2001*; *Krupic et al., 2015*; *Becu et al., 2019*). Computational models of spatial representation in rodents link this sensitivity to geometry with a postulated ability of the animal to estimate distances to surrounding walls (*Hartley et al., 2000*) or to observe panoramic visual snapshots of surrounding space (*Franz et al., 1998*; *Cheung et al., 2008*; *Sheynikhovich et al., 2009*), and rely on a wide rodent visual field (≈320°). That the width of visual field plays a role in geometric processing in humans was demonstrated in the study by ***Sturz et al.*** (***2013***), in which limiting visual field to 50° impaired performance in a geometry-dependent navigation task, compared to a control group. We thus studied whether activities of egocentric and allocentric neurons in the model encode information about the geometry of the environment and whether snapshot accumulation over time plays a role in this process.

To do this, we run the model to uniformly explore a square environment and we stored population rate vectors of the egocentric-visual and allocentric-parietal populations at successive time points during exploration. More specifically, for the egocentric population, each population vector corresponded to population activities evoked by the presentation of a single visual snapshot. In contrast, for the allocentric population, each population vector corresponded to a panoramic snapshot obtained by accumulating several successive snapshots during head rotations (see Methods). The visual information content was identical in two sets of population vectors as they were collected during the same exploration trial. Population vectors in each set can be considered as data points in a high-dimensional space of corresponding neural activities. These points are expected to belong to a two-dimensional manifold in this space, since during exploration the model animal moves in a 2D spatial plane. The analysis of the intrinsic dimensionality of both sets indeed shows that it is about 2 (*Figure 5*A,B). We then applied two different manifold visualisation techniques to see whether the shape of manifold reflects the environment shape (see Methods). We found that when applied to population vectors of the egocentric population, the structure of the manifold did not reflect the layout of the environment (*Figure 5*C). In contrast, allocentric population activities reliably preserved geometric information in the spatial organization of the manifold (*Figure 5*D). Moreover principal axes of the manifold corresponded to the principal axes of the underlying environment only for the population vectors of the allocentric population (bottom row of *Figure 5*C,D). The extraction of principal axes of an experimental space has been proposed to underlie spatial decision making in several experimental paradigms, including data from humans (*Gallistel, 1990*; *Cheng and Gallistel, 2005*; *Sturz et al., 2011*).

**Figure 5.**
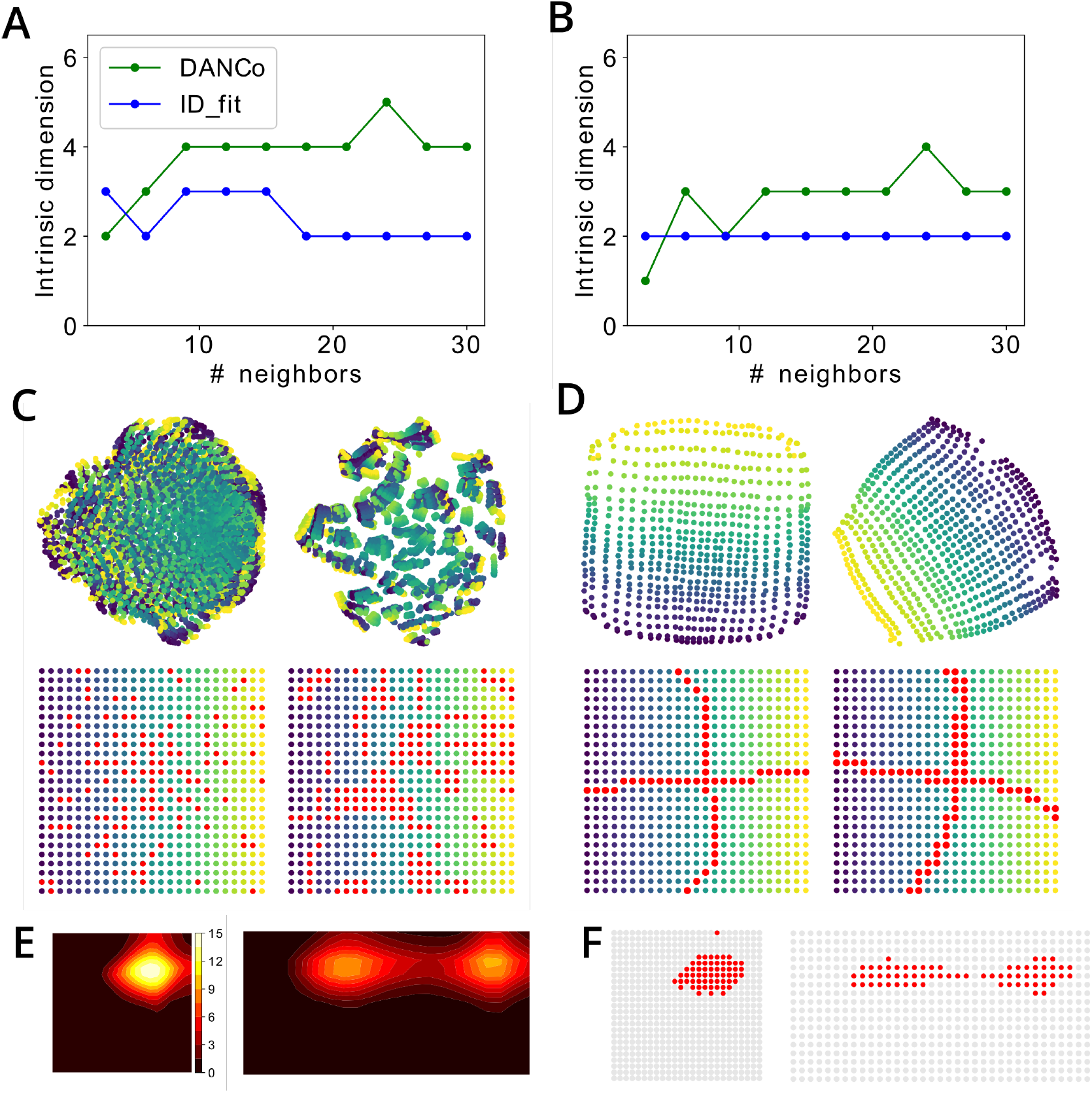
Representation of spatial relations by egocentric (occipito-parietal) and allocentric (parieto-retrosplenial) visual neurons. A,B. Estimation of intrinsic dimensionality of the set of population vectors in the egocentric (A) and allocentric (B) populations by two different state-of-the-art methods (DANCo and ID_fit). C,D. Top: Projection of the population vector manifolds onto a two-dimensional plane using Isomap (left) and t-SNE (right) algorithms. Color gradient from yellow to blue corresponds to the position at which the corresponding population vector was observed, as shown in the Bottom row. Red dots show population vectors that lie close to the principal axes of the 2D manifold of the principal space. C and D show population vectors of the egocentric and allocentric neuronal populations, respectively. E. An example of the receptive field of one hippocampal neuron after learning the environment before (left) and after (right) extension of the environment along it horizontal axis. F. For the same neuron as in E, red dots show locations in the environment where this neurons is winner in the WTA learning scheme.

STDP in the connections between the parietal and hippocampal neurons ensures that allocentric spatial views are stored in memory, while lateral inhibition in the hippocampal layer implements a competition such that different hippocampal cells become selective to different localized regions of the visuospatial manifold, which, by virtue of the coherent mapping on the real space, correspond to spatial receptive fields (*Figure 5*E). When the geometry of the environment is modified, but the memorised allocentric representation remains the same, modeled hippocampal cells express corresponding modifications of their receptive fields (*Figure 5*E,F), potentially providing a purely sensory basis for the effects of geometric manipulations observed in rats (*O’Keefe and Burgess, 1996*) and humans (*Hartley et al., 2004*). These simulations show how the egocentric-allocentric conversion and short-term memory along the modeled dorsal visual pathway can be instrumental in structuring the hippocampal input according to the geometric properties of the surrounding space that were repeatedly shown to affect human navigation (*Hermer and Spelke, 1994*; *Becu et al., 2019*).

### Visual responses of hippocampal neurons reflect learning of visual stimuli

The hippocampal memory network is thought to support a large spectrum of memory-based behaviors, and therefore its basic properties should manifest themselves in situations other than navigation. In particular, plasticity and competition, which are proposed to mediate fast hippocampal learning of visual information in our model, occur not only during navigation but also in a passive image viewing paradigm. In the next simulation inspired by the experiment of *Jutras and Buffalo* (***2010a***) we used the stationary model to learn a set of 100 novel images presented in a quick succession (see Methods) and recorded activities of modeled hippocampal neurons. In response to the presented stimuli, some neurons increased their firing rates as a result of STDP (winning neurons), while the rest of the neurons were inhibited (*Figure 6*A). Even though only a few neurons won the competition for each particular stimulus, some neurons were selective to a larger number of stimuli than others (*Figure 6*C,D). Therefore, stimulus-averaged firing rates of different neurons expressed either a decrease in the average firing rate (for neurons that were never winners), no change in the average rate (for neurons that were winners for a relatively small number of stimuli), or an increase in the average rate (for neurons that were winners for a relatively high number of stimuli, *Figure 6*B). There was a larger number of neurons that expressed decreased firing rates or no change, compared to those that increased their average rate (*Figure 6*D).

**Figure 6.**
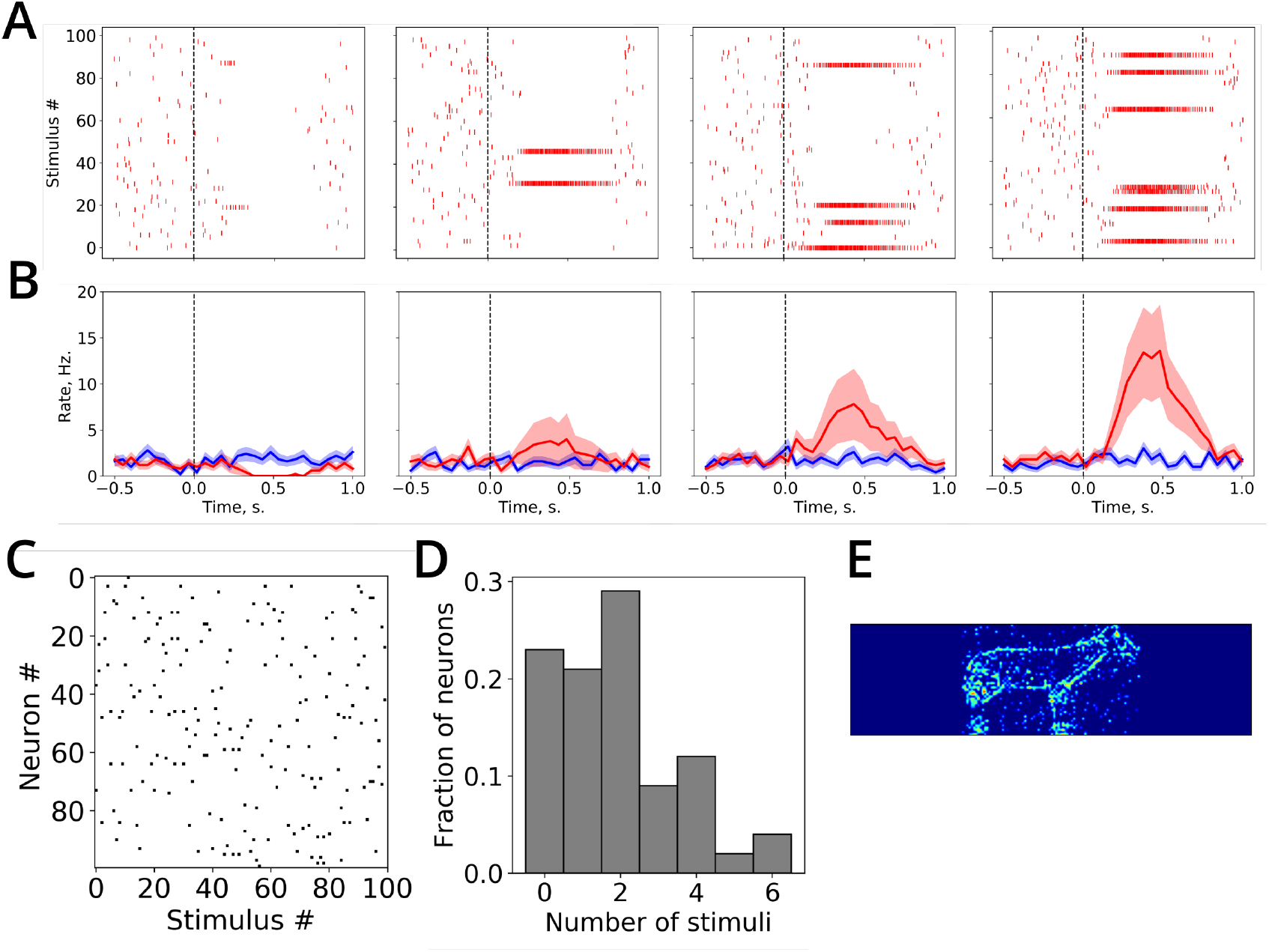
Visual responses of modeled hippocampal neurons. A. Spike raster plots for four example neurons in response to presented visual stimuli. B. Stimulus-averaged firing rates of neurons in A (mean ± SEM shown in red), compared to baseline firing rates (shown in blue). The dashed vertical line represents the stimulus onset. C. Black dots correspond to winner neurons among all other neurons (vertical axis) for each of the presented stimuli (horizontal axis). D. The histogram shows the distribution of neurons with respect to the number of stimuli for which they are winners. E. An example of the weight matrix of a hippocampal neuron after learning.

Under the assumption that a novelty-detection mechanism (assumed to reside in the hippocampus or elsewhere, but not modeled here) prevents hippocampal firing in response to a repeated stimuli, these results are in accord with the data from a number of studies showing that different subsets of recorded hippocampal neurons either decreased, showed no changes, or increased their activity in response to the presentation of a novel stimulus (*Jutras and Buffalo, 2010a*; *Rutishauser et al., 2006*). In these studies of the role of novelty in hippocampal processing, stimulus-averaged elevation of neural activity was considered as an indication of an abstract (i.e. independent of stimulus identity) novelty processing in the hippocampus. It is unclear how such an abstract representation of novelty can be reconciled with the role of the hippocampus in navigation. In contrast, our simulation results suggest that elevation or depression of stimulus-averaged firing rate in a neuron may be related to the number of stimuli for which this neuron is winner.

### Top-down hippocampal input in spatial reorientation and memory-based search

The population of the hippocampal neurons in the model represents the neural storage of (potentially highly processed) visual information aligned with an allocentric directional frame by the coordinate transformation network. In this section we show how this neural storage can affect two types of behavior: *(i)* determination of position and orientation when a disoriented monkey is placed into a familiar environment (*Gouteux et al., 2001*); and *(ii)* memory-guided visual target search in an image viewing paradigm (*Fiehler et al., 2014*). While these two tasks may seem unrelated, we propose that the same neural process, namely a reorientation of the head-direction network based on the comparison between the newly obtained visual information and the contents of the hippocampal allocentric storage, underlies behavioral decisions in these tasks.

#### Spatial reorientation

In a series of reorientation experiments with monkeys, *Gouteux et al.* (***2001***) have shown that these animals relied on both the geometric information (given by the three-dimensional layout of the rectangular experimental space) and non-geometric cues (e.g., landmark objects placed near the walls or corners of the recording chamber). The authors paid specific attention to the influence of landmark size on reorientation behavior. When small objects were placed near one of the walls or in the corners of the room, the monkeys did not use these cues to reorient, and their search pattern was determined based only on the geometric information. Importantly, this was not because the monkeys did not notice the landmarks, since they performed exploratory actions towards them (looked at or touched them). Once the landmark size was increased however, the monkeys successfully used them for reorientation independently of their location and number (i.e. when they were present).

To simulate these data, we tested the model in four reorientation tasks in a virtual three-dimensional rectangular room. In these tasks, either no landmark cues were present in the room, or one visual landmark of three different sizes was placed in the middle of one of the walls (*Figure 7*A). Each task comprised an exploration phase, during which the model randomly explored the environment, and a reorientation phase. In the reorientation phase the model was initialized with a random heading direction and placed back into the environment learned during the exploration phase at a random location. The performance of the model was assessed from the accuracy of reorientation: we assume that the animal will navigate to the correct corner if it has correctly estimated its initial heading, whereas it will make a navigation error if the reorientation error is high.

**Figure 7.**
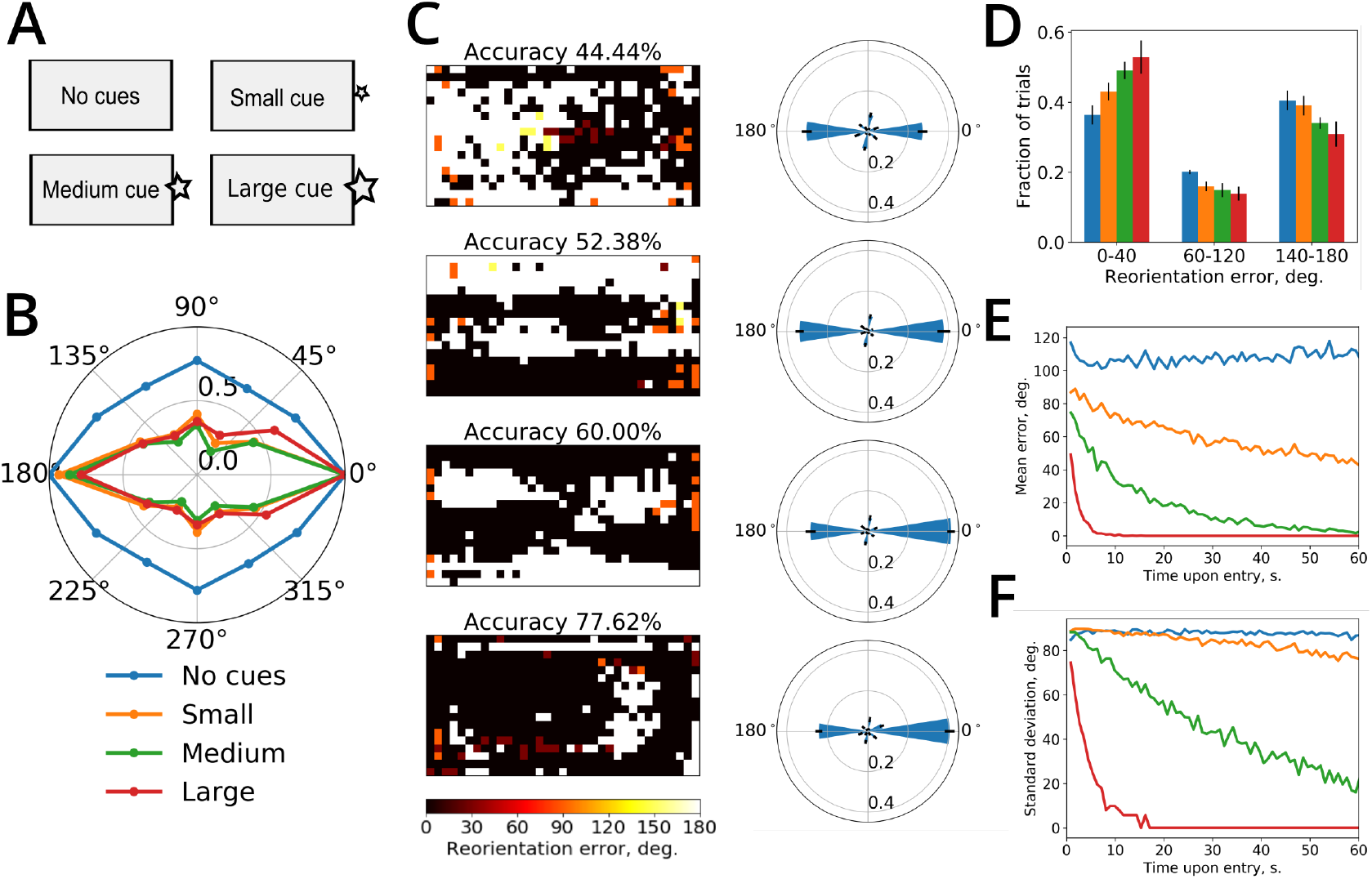
Simulation of the reorientation experiment. A. The experimental environment was a rectangular room (represented by the gray rectangles). The same reorientation simulation was run in four conditions: no visual cues apart from walls of the room, or 1 visual cue at three different sizes (small, medium, large). B. Polar plot of the mean activity of the reorientation network when the simulated animal was placed in various locations in the room. Dots mark the preferred locations of the *N*_re_ reorientation neurons. Colors from blue to red represent 4 experimental conditions. C. Rows from top to bottom correspond to experimental conditions as in A. Left: Reorientation maps show, for each location in the room, the reorientation error committed by the model after seeing only the first visual snapshot from that location (at a randomly chosen head orientation). The pixel color from black to white codes for the absolute value of the reorientation error from 0 to *π*. Right: polar histograms of reorientation errors (±SD), averaged over 9 random orientations at each location. D. Bar plot shows the distribution of the absolute reorientation errors (±SD) among the approximately correct orientation (0-40°), rotational error (140-180°) and other directions. E,F. Reorientation error mean (E) and its standard deviation (D) when progressively more snapshots were used for reorientation. Color code for D,E,F as shown in B.

Once the information from the initial view reached the hippocampus upon the reentry to the environment, the activity peak of the reorientation network signalled the orientation error (*Figure 7*B). This error represented the discrepancy between the initial heading direction and the heading direction most consistent with the allocentric information stored in the projections from the place cells to the reorientation network. The asymmetric shape of the polar plot reflects the influence of the environment’s geometric layout on reorientation: for the no-cue condition, the network activity peaked at the correct (0°) and its rotationally opposite (180°) orientations with an identical average amplitude. When the visual cue was present, its size determined the difference between the activity peaks. Therefore, when reorientation was performed from different locations in the environment (based only on the first view taken), the accuracy, measured as the percentage of locations with a correctly determined orientation, was about 44% in the no-cue condition and raised to about 78% in the large-cue condition (*Figure 7*C, left column). These reorientation maps show that depending on the position of the orienting cue in the room and the view taken, some locations in the environment provide better visual information for reorientation than others (shown by white areas in the maps). The histograms of orientation errors (*Figure 7*C, right column, and *Figure 7*D) show that, on average, a larger visual landmark provides a much better reorienting cue than a small one, for which a similar number of correct decisions and rotational errors was observed (*Figure 7*D). This is due to the fact that orientation is determined essentially by comparing the egocentric view from the initial position with allocentric views stored in synaptic memory, without any explicit landmark identification process. Therefore, influence of small visual cues becomes negligible with respect to gross visual features of the surrounding space (corners, outline of the walls, etc.). These results are consistent with the hypothesis that reorientation is a fast, bottom-up process based on low-level visual information (*Sheynikhovich et al., 2009*). Learning landmark identities and their spatial relation to goals can be added by subsequent learning, but may not be taken into account unless they are sufficiently salient compared to geometric cues present in the environment (*Cheng, 1986*).

So far the reorientation performance was assessed based only on the first view taken. It is likely to increase if the animal is allowed to accumulate visual information from successive views taken in the same location at different orientations or at different locations, e.g. during initial movements through the environment. This is what happens in the model, since increasing the number of snapshots that are used for reorientation improved its accuracy (*Figure 7*E,F). In this case we placed the simulated animal at 60 successive positions, while at each position the animal rotated its head to obtain a corresponding panoramic view. The activity of the reorientation network was calculated as a sum of its activities after each successive view. When a large cue was present, the simulated animal obtained an accurate orientation estimate after visiting about 10 successive locations. In contrast, the mean error and standard deviation of reorientation were decreasing much slower for smaller sized landmarks. Thus, our model describes a neural mechanism for spatial reorientation which relies on an allocentric visual information stored in the hippocampal network. This allocentric information feeds into a head-direction-like network, assumed to reside in the retrosplenial cortex, that signals reorientation error and affect the sense of direction via its input to the head-direction system if the brain (*Taube, 2007*). In addition to providing a mechanistic basis for the reorientation process, which is a necessary part of navigational behavior and whose existence is assumed (either implicitly or explicitly) in a number of computational models of navigation, this model proposes how reorientation can be performed continuously, i.e. during ongoing spatial behavior.

#### Memory-based visual search

Finally, to illustrate a potential role of the stored hippocampal representation in memory-based visual tasks, we simulated the study of ***Fiehler et al.*** (***2014***). In this study, head-fixed human subjects remembered a visual scene with 6 objects on a table, presented on a computer screen (*Figure 8*A, top). This encoding phase was followed by 2-s. delay (uniform gray image), and then the subjects were presented with a modified scene in which one of the objects was missing (the target object) and either 1, 3 or 5 other objects displaced horizontally (*Figure 8*A, bottom). The subjects were required to point to the remembered location of the missing object. If the subjects had used only an egocentric information (i.e. remembered object position with respect to the head), then their performance would have been independent from the displaced objects. The results of this experiment demonstrated in contrast that pointing performance was influenced by the non-target objects, such that shifting a higher number of them induced a larger pointing error. Even though the pointing error was always made in the direction of the object displacement in the image, the size of the error only partially accounted for the veridical displacement of the objects. These data suggest that human subjects combine allocentric (i.e. based on the information from the environment, in this case represented by the visual features associates with displaced objects) and egocentric (i.e. based on the memory of an egocentric location of the target object) information during memory-based search (*Fiehler et al., 2014*). The neural mechanism of this allocentric correction of the egocentric memory is unknown.

**Figure 8.**
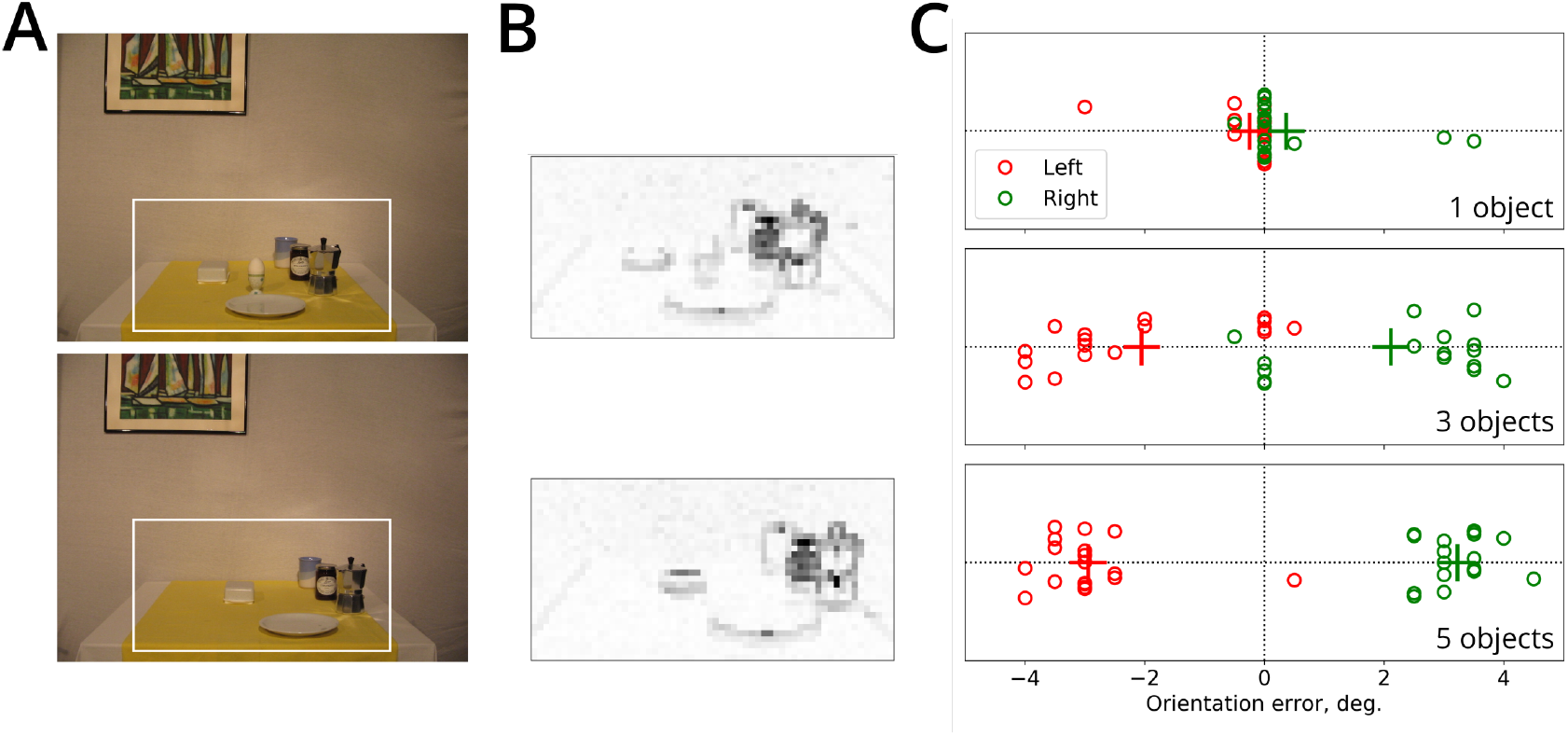
A. An example of the remembered (top) and test (bottom) images. In this example, the target object is the egg and 5 non-target objects were shifted to the right in the test image, compared to the encoded image. The white rectangle denotes the part of the image that was provided as input to the network. It corresponds to the part of the image most fixated by the subjects in the experiment. B. Mean firing rates of the egocentric neurons in the model for the encoded and test images shown in A. C. Orientation errors induced in the model by the presentation of the test images with 1 (top), 3 (middle) and 5 (bottom) displaced objects. Horizontal position of each dot corresponds to the maximal activity peak of the reorientation network. Different dots represent different sets of objects in the image dataset. Leftward and rightward displacements are shown in red and green, respectively. Crosses mark the mean displacement value per group. Random jitter along the vertical axis is added for clarity.

We hypothesized that the influence of allocentric image information observed in this experiment arises as a result of a slight misorientation of the head direction network due to the apparent shift of visual features caused by the object displacement in the attended area of the image. In order to demonstrate this effect, we first presented to the model an image of a control scene with all 6 objects (see *Figure 8*A, top, for an example). We used, with permission, the same image dataset that was used in the experimental study. As an input to the network we only used the part of the image near the objects, because in the experiment it was fixated most of the time and because of the evidence that displacement of objects outside of this area had no influence on reaching performance (*Fiehler et al., 2014*). The network converted the visual input of the egocentric layer (*Figure 8*B) to an allocentric representation according to the actual head direction (set to 0°), which was stored in the synapses between the parieto-retrosplenial output cells and hippocampal cells as before. In this simulation we ignored competition effects, since it was not required to remember multiple images. Second, after the first scene was learned, an image of the scene with one object missing and either 1, 3 or 5 objects displaced (see *Figure 8*B, bottom) was presented to the model. The orientation error caused by the object displacement can then be read directly from the activity of the reorientation network (*Figure 8*C). As in the experiment, the number of displaced objects affected the amount of allocentric correction. Since in the test images the displaced objects correspond only to a subset of all visual features, the mean correction only partially account for the object displacement. Thus, as in the case of spatial reorientation, the influence of the allocentric information (in this case represented by low-level features of the presented image) is caused by the comparison between the stored allocentric and incoming allocentric views, and the resulting activity of the reorientation network that calibrates the head direction signal.

## Discussion

The presented model focuses on the dorsal visual pathway for information processing, generally thought to provide contextual or “where” information to memory structures in the MTL, by contrast to the ventral pathway mediating the processing of object/item representations or “what” information (*Goodale and Milner, 1992*; *Kravitz et al., 2011*). The two pathways converge at the hippocampus where both types of information are combined to form episodic memories. In both spatial (e.g. spontaneous novelty exploration) and non-spatial (recollection/familiarity) experimental paradigms the dorsal pathway has been implicated in the encoding of contextual information (e.g. the scene or location where an item was observed) and not in remembering the object identity (see *Eichenbaum et al., 2007*, for review). These proposals go in line with general properties of neural activities observed in brain areas along the dorsal pathway such as PHC and RSC. In particular, fMRI studies show that these areas are activated by scene processing, with PHC responding equally strongly to images of spatial layouts with or without objects (*Epstein and Kanwisher, 1998*; *Epstein, 2008*). RSC was shown to be more strongly implicated in recollection than familiarity (*Epstein, 2008*) and is proposed to play a specific role in encoding spatial and directional characteristic of landmarks and their stability *independent of their identity* (*Mitchell et al., 2018*). The central claim of our model is the existence of an allocentric representation of surrounding visual space in topographic visual coordinates, as emerging experimental evidence suggests (*Melcher and Morrone, 2015*; *Park and Chun, 2009*; *Robertson et al., 2016*; *Julian et al., 2018*). We propose that such a representation can be accumulated from successive views during head rotations independently from mechanisms ensuring transsaccadic memory and apparent visual stability (*Wurtz, 2008*), and provide a neural interface between visual and mnemonic structures during spatial orientation. We further show how such a representation can support memory-based visuospatial behaviors such as spatial reorientation and memory-based visual search. We believe that such a representation is necessary to explain experimental data demonstrating the role of memory in visual behavior in a number of experimental paradigms (*Chun and Nakayama, 2000*; *Tatler and Land, 2011*; *Robertson et al., 2016*; *Laeng et al., 2014*). These issues and related predictions from the model are further discussed below.

### Contextual memory representations

In the present work, the selectivity to scenes and spatial layouts, as opposed to objects, is modeled simply as sensitivity to integrated visual ‘snapshots’ (i.e. the contents of the animal’s visual field acquired across multiple fixations and accompanying head movements). The integration time scale is proposed to be defined by the time constant of the short-term memory processes necessary for snapshot accumulation. In our model we chose to represent the contents of a view by a retinotopic-like grid of orientation-sensitive filters for simplicity, but the coordinate-transformation circuit and the rest of the model are in fact agnostic about the nature of features that enter the transformation network as long as they are represented in retinotopic-like head-fixed frame and take up the whole visual field. These could for example combine low-frequency content of input views (*Schyns and Oliva, 1994*; *Kauffmann et al., 2015*), depth cues (*Lee et al., 2012*; *Raudies and Hasselmo, 2012*), salience maps or other perceived features computed within the occipito-parietal network. Object processing is assumed to be done in parallel in the ventral stream, since object representations are generally considered to be view-independent and assume translation invariance over the visual field (*Serre et al., 2005*). The relative (i.e. size dependent) sensitivity to objects in our model (see “Spatial reorientation”) arises from the fact that large, distal and stable objects (or landmarks) that make up a large portion of a view are considered as part of the layout, and not as identified objects/landmarks. In contrast, relatively small objects, landmarks, or a high-frequency contents of other small localized portions of a view contribute only weakly to the overall visual representation. Indeed, they are often overshadowed by gross visual features present in views, such as corners, walls, and other large-scale visual structures during comparison of new and remembered view-based representations (*Cheng, 1986*; *Schyns and Oliva, 1994*).

Our model can thus be considered as a model of encoding of contextual information, as opposed to object-related one, and the notion of context is well defined: it is the visual information provided by topographically-organized features present in a set of views and stored in memory after the acquisition phase of a task. This notion of context to can be extended to a non-spatial setting (see “Memory-based visual search”): topographically-organized image features present in attended part of the screen and stored in memory provide contextual information with respect to any object-related information (such as object identities) stored from the scene. The importance of mnemonic component in contextual spatial memory is supported by the evidence for ‘priming of pop-out’ and ‘contextual cueing’ (*Chun and Nakayama, 2000*). In the latter paradigm, for example, subjects performed visual search of a letter T among rotate Ls. Even though the subjects did not know that the the spatial layout of presented letters was predictive of target spatial location, their performance was better in previously observed contexts than in novel ones. That is, subjects implicitly remembered previously presented scenes and used them to drive visual search. In the absence of reliable object-related information (such as spatial scene without conspicuous landmarks), contextual information can be used to drive attention and visual behavior. The important piece of information that is present in topographic representation of a scene, but is absent in object-related memory, is spatial location. Indeed, one can assign position information within the topographic representation of visual space (with respect to an allocentric directional frame, or with respect to other visual features). Therefore, (allocentric) contextual representations can serve as a basis for remembering spatial and directional characteristics of objects or landmarks independent of their identity. Spatial locations in such a contextual representation can serve as “place holders” for specific object/landmark information extracted and stored in the ventral visual stream, or as “pointers” to this information (*Cavanagh et al., 2010*). Such a notion of contextual information is well in line with a proposed role of the PHC and RSC in landmark processing (*Epstein, 2008*; *Mitchell et al., 2018*).

### Spatiotopic representations in the brain

While the existence of view-based representations in human spatial memory is well established (*Shelton and McNamara, 1997*; *Diwadkar and McNamara, 1997*; *Christou and Bülthoff, 1999*; *Garsoffky et al., 2002*; *Burgess, 2006*), the existence of a spatiotopic representation of the surrounding visual space is rather controversial. The existence of spatiotopic representations in the brain has traditionally been discussed in the context of apparent stability of the visual world despite saccadic eye movements that abruptly change retinal image 2-3 times per second on average (*Pylyshyn, 1989*; *O’Regan, 1992*; *Melcher, 2005*; *Wurtz, 2008*; *Cavanagh et al., 2010*). A discovery of visual remapping (sometimes referred to as ‘shifting receptive fields’, *Duhamel et al., 1992*) and elucidation of neural mechanisms responsible for it (*Wurtz, 2008*) provides a rather strong evidence for “a minimalist theory of visual stability – keep track of the location of currently attended items and nothing else is required” (*Cavanagh et al., 2010*). However, substantial literature on spatiotopic adaptation effects suggests that spatiotopic representations may coexist with retinotopic ones but can only be observed when some necessary conditions are met (*Melcher and Morrone, 2015*). In other words, the conclusion that spatiotopic representations are not useful to explain apparent visual stability does not mean that they are not required for other functions. The retinotopic vs spatiotopic debate in the context of visual stability resembles egocentric vs allocentric debate in the context of spatial cognition (*Wang and Spelke, 2002*; *Burgess, 2006*). Proponents of the purely egocentric view propose that egocentric updating of several locations of interest is all that is needed for spatial navigation, which can be true for the case when only a few locations have to be tracked (*Wang, 2017*). However, when the complexity of the task increases, an allocentric representation of space represents a much more efficient way of storing spatial information (*Burgess, 2006*). In an analogous manner, the proposed spatiotopic representation of visual space provides an efficient way of storing and accessing memory about spatial scenes building blocks of allocentric spatial representations.

On the basis of our results we propose that three aspects are important for future experimental validation of the model. First, the nature of the stimulus and temporal aspects of stimulus presentation can be important for observing spatiotopic effects. In terms of our model, while visual adaptation phenomena in retinotopic coordinates are assumed to occur in purely retinotopic areas of the brain (i.e. in the occipito-parietal network, of which only craniotopic output component is explicitly modeled), the spatiotopic effects can only take place after the transformation network, i.e. deep within the hierarchy of the parietal-retrosplenial circuit. The model thus predicts that spatiotopic effects should require longer presentation times to be observed and they should be observed for moderately complex stimuli, such as those encoded by neurons in the higher-level visual hierarchy projecting to parietal areas (i.e. before a ‘branch point’ to ventral representations of complex objects and faces). If true, these spatiotopic effects should be preserved during head rotations, even if the location of the stimulus temporarily leaves the visual field (for the time shorter than the short-term memory time scale). Inactivation of head direction cells in the RCS is predicted to prevent spatiotopic effects during head rotations and preserve retinotopic ones.

Second, while retinotopic effects are tightly linked with attention, spatiotopic representations are intrinsically connected to memory, so that mnemonic component of the task can be important for the spatiotopic effects to be observed. Moreover, multiple lines of evidence suggest that the actual contents of the visual field overrides any memory of these contents, and, potentially, any related spatiotopic effects. For example, *Oliva et al., 2004* tested visual and memory search strategies in panoramic scenes. When searching for objects a priori known to be present in the visual display, subjects used inefficient visual search strategy over hundreds of presentations of the same scene, even though a memorized contents of the scene could have been used more efficiently. However, when objects were located in the scene outside of the current visual field, memory of the scene was readily used. In another study testing repetition-suppression (RS) effects at the level of the PHC using fMRI, *Golomb et al., 2011* observed RS when subjects performed saccades through a stationary scene, suggesting spatiotopic processing. However, the same RS effect was observed when eyes remained stationary, but the scene moved as if the same saccades were made, suggesting that spatiotopic memory effects were discarded when the apparent visual field was the same. These and similar data show that information in memory (which is proposed to be stored in an allocentric spatiotopic frame) is discarded when the actual information in the visual field (i.e. in the retinotopic frame) can be used instead. These considerations suggest that spatiotopic effects should be more easily observed in tasks that require recall of visual information from memory, as opposed to tasks that could be solved using information present in the screen.

The third aspect concerns the nature of the task to be solved by the subject. As briefly mentioned in the introduction, the dorsal stream network can be subdivided into three distinct major pathways, mediating executive (working memory), sensorimotor (visually-guided action) and spatial navigation functions (*Kravitz et al., 2011*). It is thus reasonable to suggest that different pathways rely on suitable reference frames. While less clear for the executive pathway, visually-guided action can be assumed to rely mostly on egocentric (eye- or body-based) frame, whereas spatial navigation is likely to use allocentric reference frames. Therefore, using spatial navigation context in visual tasks can bias the visual system to express spatiotopic vs retinotopic processing.

Whereas the allocentric representation in our model is purely visual, the possibility that it could be multisensory can not be excluded (*Newell et al., 2005*). *Loomis et al., 2013* defined a similar representation of surrounding 3D space as a “spatial image” with the following properties: *(i)* it can be updated during movement with the eyes closed; *(ii)* it exists in all directions; *(iii)* the information from all sensory modalities converge onto a common, “amodal”, spatial image. While our model is directly consistent with the second property, the third one can be implemented by converting spatial locations of egocentric sensory signals at different modalities (e.g. haptic or auditory) into the common allocentric framework. These locations (or placeholders) can then be linked to the detailed representations of sensory experience in sensory-specific areas of the cortex, similarly to the putative links between landmark locations and their high-frequency contents discussed above. The first property can in principle be assured by backward projections from the hippocampus to the allocentric layer (not included in the model) by a mechanism previously proposed to support spatial imagery (*Byrne et al., 2007*), it is not clear however how such backprojections can be learned in a biologically plausible neuronal network.

One obvious candidate for the potential biological locus of the panoramic visual representation is the PPC, since spatiotopic neuronal receptive fields were observed in this area (*Galletti et al., 1993*; *Snyder et al., 1998*; *Fairhall et al., 2017*). Another potential candidate is the RSC that was recently proposed to mediate a memory of panoramic visual scenes (*Park and Chun, 2009*; *Robertson et al., 2016*). The parahippocampal place area, a region of the PHC known to participate in scene processing (*Epstein and Kanwisher, 1998*), is unlikely to mediate such a representation, as it did not show RS effects during presentation of overlapping views taken from the same panoramic scene (*Park and Chun, 2009*).

### Novelty encoding in the hippocampus

Single cell recordings in monkey and human hippocampus provided evidence for novelty and familiarity encoding in this area: ‘novelty-detecting’ neurons strongly increase or decrease firing rates in response to the appearance of visual stimuli the monkey have never seen before, while ‘familiarity-detecting’ neurons significantly modify their firing rates only in response to previously encountered stimuli (*Rutishauser et al., 2006*; *Jutras and Buffalo, 2010a*). In these studies sensitivity to novelty was assessed by averaging neuronal firing rates across multiple stimuli. Elevated average rates of some neurons in response to novel stimuli were taken as evidence of a general (i.e. stimulus-unspecific) novelty signal in the hippocampus. Our results argue against the concept of general novelty signal, since in the model it is an artifact of stimulus-averaging: an increase or a decrease of the average rate of a neuron depends on the number of stimuli to which the neuron is sensitive (i.e. for which the neuron is ‘winner’, *Figure 6*B). The competitive learning account also provides an explanation of why the majority of novelty-detecting hippocampal neurons decreased their firing rates in the study by *Jutras and Buffalo, 2010a*: each stimulus is encoded by a small number of neurons that inhibit the rest of the network. Our results thus go in line with the proposals linking novelty signals with competitive encoding of stimuli (*Viskontas et al., 2006*). While our simple model of spatial coding accounts for an elevated or a depressed firing rate of a neuron in response to a novel stimulus during encoding (that presumably occurs in the EC or CA1), it can not explain neither familiarity response (i.e. when a neuron changes rate only upon the second presentation of a stimulus) nor repetition suppression (i.e. the absence of rate changes when a previously observed stimulus is presented) that are related to stimulus retrieval from memory. Taking into account these phenomena will require the addition of a memory structure, such as CA3, which is out of the scope of the present work (see *Hasselmo et al., 1995*; *Norman and O’Reilly, 2003*, for related models).

A recent discovery of ‘concept cells’ in the human MTL was primarily based on object coding properties of these cells mediated by ventral visual pathway (*Quiroga, 2012*). These cells are thought to arise as a result of sparse coding in hierarchically organized MTL processing. Our model, similarly to many previous computational models of spatial coding (e.g. *Hartley et al., 2000*; *Arleo et al., 2004*; *Byrne et al., 2007*; *Sheynikhovich et al., 2009*), proposes that place cells arise via similar competitive processing but based on the visual inputs lacking detailed object-related information. It extends these previous models by showing how such visual inputs can be formed in the primate dorsal visual path networks and therefore contributes to the understanding of how episodic memory is formed based on the conjunction of place- and object-related information by the hippocampal networks (*Miller et al., 2013*; *Ison et al., 2015*).

### Neural mechanisms of spatial reorientation

In the present work, reorientation (i.e. recovery of location and heading after disorientation) is proposed to be performed by comparing a currently perceived view (or a collection of recently observed views) with the total allocentric sensory information stored in the hippocampal network during training (see also *Sheynikhovich et al., 2009*). This comparison results in the activation of hippocampal cells corresponding to locations with similar allocentric features, resulting in a distributed representation of spatial location. This representation can then be used drive navigation based on reward-based learning. This account is different from a classical “view-matching” approach in which the current view is matched only with one remembered reference view (e.g. the view taken at the goal location), and the animal is assumed to move so as to minimize the difference between these views without building an intermediate representation (*Franz et al., 1998*; *Stürzl et al., 2008*). A set of all memorized (allocentric) views stores much more information than just a reference view. As we show in the Results (*Figure 5*), the collection of stored allocentric views can be considered as belonging to a two-dimensional manifold that is mapped to the actual space and it can therefore define a coordinate frame with respect to which spatial locations can be coded. Moreover, if the mapping is accurate, principal axes of the manifold correspond to the principal axes of the actual space. Recent work in rodents has shown that hippocampal spatial representations can be built after complete lesions of the medial entorhinal cortex that is thought to encode self-motion and boundary-related information in grid cells and border cells (*Schlesiger et al., 2018*). How the remaining inputs can result in reorganization of spatial maps is an open question. Here we show that a collection of stored allocentric views can be a basis for creating such representations (see also *Li et al., 2020*). A number of studies in rodents and humans suggested the sensitivity of spatial memory to intrinsic axes of observed sets of spatially organized stimuli (*Gallistel, 1990*; *Cheng and Gallistel, 2005*; *Sturz et al., 2011*; *Marchette et al., 2011*) and the neurophysiological basis for this sensitivity is not clear. Our results suggest that information about these intrinsic axes is implicitly stored in the set of stored allocentric views. Application of view-matching approaches to explain human navigation is often criticized based on observations suggesting the use of depth information and not image-matching (*Lee et al., 2012*). However, if distances to surrounding objects/boundaries are represented in a retinotopic frame, they can be thought of as images (or ‘depth maps’ in computer graphics). Thus, any image-matching algorithm (including ours) can be converted to a distance-matching one by a suitable modification of the input.

In contrast to a previous model of reorientation that relied on the presence of conspicuous landmarks linked directly to head direction cells (*Bicanski and Burgess, 2016*, ***2018***) our model proposes a mechanism of fast bottom-up view-based reorientation of the head-direction signal that does not require landmarks (polarized geometry is sufficient). This is in accord with a number of reorientation studies mentioned earlier suggesting that this neural process is independent from landmark identities and can be performed in the absence of point-like landmarks. The mechanism we use relies on weight sharing and as such is not, at its present implementation, biologically realistic. We note here that the concept of weight sharing has been critical for recent successes of brain-inspired neural networks and is widely used in models of biological networks of visual processing (e.g. *Serre et al., 2005*; *Masquelier and Thorpe, 2007*; *Bartunov et al., 2018*). We see two possibilities of how the reorientation process can be implemented in a more realistic network. One possible implementation would require a mental rotation of the stored allocentric representations, while freezing the actual egocentric view in the input layer. A match is found when the maximal activity of the reorientation network reaches a threshold. We have not attempted to implement such a network due to its complexity, but we conjecture that the complexity of this computation should be experimentally detectable by a prolonged reaction time compared to a control setting. A second possibility is that the comparison of a currently perceived view with hippocampal memory contents is performed with the help of ongoing behavior: a disoriented subject could for instance actively rotate its head or body until the match is found. A prediction in that case is that the subject’s orientation just before taking the decision should predict future behavior. For example during hidden goal navigation in a rectangular room, the head orientation before movement initiation should be predictive about a correct behavior or a rotational error.

### Conclusion

To summarize, the model presented in this work explored the nature of visual representations in the parietal-medial temporal pathway for visuospatial processing and contributed to the open question of the link between visual and memory structures in primates. It proposes that a single, potentially multisensory, representation of surrounding environment is constructed by time-integrated sensory snapshots. This putative representation provides a coordinate space within which positions of localized sensory events can be encoded and that can represent visual locations outside the current field of view. This model thus provides a conceptual framework for linking oculomotor behavior, visual and spatial memory.

